# Colonization, translocation, and evolution of opportunistic pathogens during healthcare-associated infections

**DOI:** 10.1101/2025.10.23.683921

**Authors:** Martin Fenk, Alexandra Hrdina, James B. Winans, Meike Soerensen, Laura Ostertag, Elina Coquery, Fatoumata Sow, Sirak Petros, Catalina-Suzana Stingu, Norman Lippmann, Carey D. Nadell, Igor Iatsenko, Bastian Pasieka, Felix M. Key

## Abstract

Many commensal bacteria that peacefully reside in the human microbiome are also able to cause acute opportunistic infections. Emerging evidence suggests that within-host evolution contributes to infection, but the genetic mechanisms facilitating the progression of opportunistic pathogens from carriage to acute infection remain unknown. Here, we prospectively collected native samples from four microbiome niches of 13 critically ill patients to assess the evolutionary dynamics leading up to infection. Among three patients we have observed eleven healthcare-associated infections (HAI) caused by nine pathogen species. Leveraging a culture-based approach, we demonstrate that the microbiome is frequently (73%) colonized by the pathogen lineage already before or at the time of diagnosis. Moreover, we identify a short-lived, non-synonymous mutation (F126L) within the fimbriae regulator gene *fimZ* of *Enterobacter hormaechei*, first detectable within the gut and subsequently associated with HAI before becoming replaced by body-wide sweeps of independent treatment-associated mutations. Despite *fimZ* [F126L] being globally undetected, we can show *in vitro* and *in vivo* that the F126L mutation leads to elevated biofilm formation, cell adhesion and virulence, suggesting a role during HAI. Our work highlights the power of prospective, population-wide investigation of pathogens to elucidate rapid evolution linked to disease.

## Introduction

Although human commensals have co-evolved with their host ^1^, many bacterial species that peacefully colonize a microbiome niche are also capable of causing acute infections. These bacteria, known as opportunistic pathogens ^2^, pose a significant threat, particularly to critically ill or immunocompromised, hospitalized patients, causing millions of healthcare-associated infections (HAIs) annually ^3–5^. The mechanisms leading to HAI are complex including the underlying disease and its severity, comorbidities, use of organ support therapy, surgery and presence of invasive devices, leading to an overall incomplete understanding of why some patients suffer from HAIs and others not. Interestingly, analyses of opportunistic pathogens from patients during infection have provided evidence for within-host translocation, as well as mutations that are associated with pathogenicity and treatment resistance which possibly contribute to disease initiation ^6–20^. While these findings support the concept that opportunistic infections aided by evolutionary mechanisms emerge from the microbiota within the patients, the mechanisms leading to opportunistic infections within human bodies have not been studied.

Theoretical, experimental, and *in vivo* evidence raises the possibility that adaptive within-host evolution can promote opportunistic infections. The rate at which bacteria accumulate mutations over time has long been underestimated. We now appreciate that billions of *de novo* mutations are generated within a human microbiome daily - an enormous adaptive potential for opportunistic pathogens residing in our body ^21^. The genetic processes contributing to within-host translocation and adaptation during the transition from carriage to acute infection of an opportunistic pathogen are poorly understood, despite intriguing evidence for their importance. For example, pervasive adaptation of microbial populations has been shown in experimental evolution ^22,23^, and its importance during infection onset in animal models has been identified ^24,25^. Interestingly, tracing asymptomatically colonizing *Staphylococcus aureus* in a single patient over one year demonstrated that the progression to fatal blood-stream infection was associated with *de novo* mutations that truncated pathogenicity-associated transcriptional regulators ^26^. Together these studies point towards a complex realm of evolution within human microbiomes contributing to opportunistic infections.

Detecting disease-promoting mutations is challenging, because such mutations may be absent or rare within microbiomes prior to disease. In addition, the plasticity of microbial genomes and the pleiotropic nature of some genes ^27^ might render vastly different genotypes adaptive for disease-associated traits such as translocation, invasion, and virulence. As such, mutations conferring a context-specific advantage may be highly individualized and rare on a global level. Moreover, adaptive mutations may be short-lived during disease and become extinct quickly if they are associated with a fitness cost, blocked from onward transmission or outcompeted by other more fit genotypes ^28^. Classic metagenomics of patient material lacks the resolution necessary to trace the dynamics of individual adaptive mutations within the genetic diversity of a given species population colonizing the patient’s body. Instead a population-wide, whole-genome approach of the pathogen species coupled with prospective sampling of patients’ microbiome niches is necessary to illuminate the progression from asymptomatic carriage to infection.

Here, we investigate the microbiome and clinical isolates sampled longitudinally from prospectively enrolled critically ill patients to retrace the cryptic evolutionary, temporal, and spatial path promoting HAI. We have observed eleven HAIs among three patients, and provide evidence for microbiome colonization preceding infection, and once even before hospitalization. During an *Enterobacter hormaechei* infection, we have found a non-synonymous mutation in the fimbriae regulator *fimZ* first detected in the gut microbiome becoming associated with pneumonia but subsequently swiftly replaced through independent genotypes linked to treatment resistance. We show that isolates carrying the *fimZ* mutation have increased biofilm formation, and lung epithelial cell adhesion *in vitro* and virulence *in vivo*. At the same time, the mutation is absent in a global genome dataset of clinical, respiratory-associated *E. hormaechei* genomes suggesting it is a rare disease-associated mutation.

## Results

### Prospective, body-wide reconstruction of pathogen diversity in healthcare-associated infection

The combination of prospective native sampling at four microbiome niches coupled with clinical isolate collection identifies the pathogen across the patient’s body. At the participating intensive care unit, clinicians screened 174 critically ill patients over a period of 3.5 months and identified 25 (14.4%) eligible patients, of whom 13 (52%) were included in this study (**Figure S1**, Methods). For each of those patients, weekly viable samples of four different microbiome niches (oral, nasal, rectal, and skin) were collected and their clinical progression and treatment were recorded. Among this cohort, three patients (patient 7, 10 and 21; 23%) developed multiple, often recurrent, episodes of HAIs caused by two to five different pathogenic species (ten respiratory tract infections and one blood-stream infection [BSI]) during hospitalization (**Figure 1b**, **Table S1, Table S2**). Upon culture-based identification of the pathogen species, up to 10 viable isolates of the pathogen species per clinical sample were collected and stored (**Figure 1b**), which due to logistical challenges during the COVID19 pandemic represents only a subset of all culture-positive HAI detections among the three patients (**Figure S2**, **Table S2**). Next, we aimed to identify the bacterial pathogen within the patients’ microbiome via targeted culturing of the prospectively collected viable nasal, oral, rectal, and skin swabs. Apart from both *Pseudomonas aeruginosa* HAI, we successfully cultured each pathogen species within at least one of the respective patients’ microbiome samples among the nine remaining HAI. Notably, in those nine pathogen-patient pairs we were able to culture the pathogen species prior to or within 6 h of HAI onset from at least one microbiome sample (**Figure 1b**, Methods).

**Figure 1:**
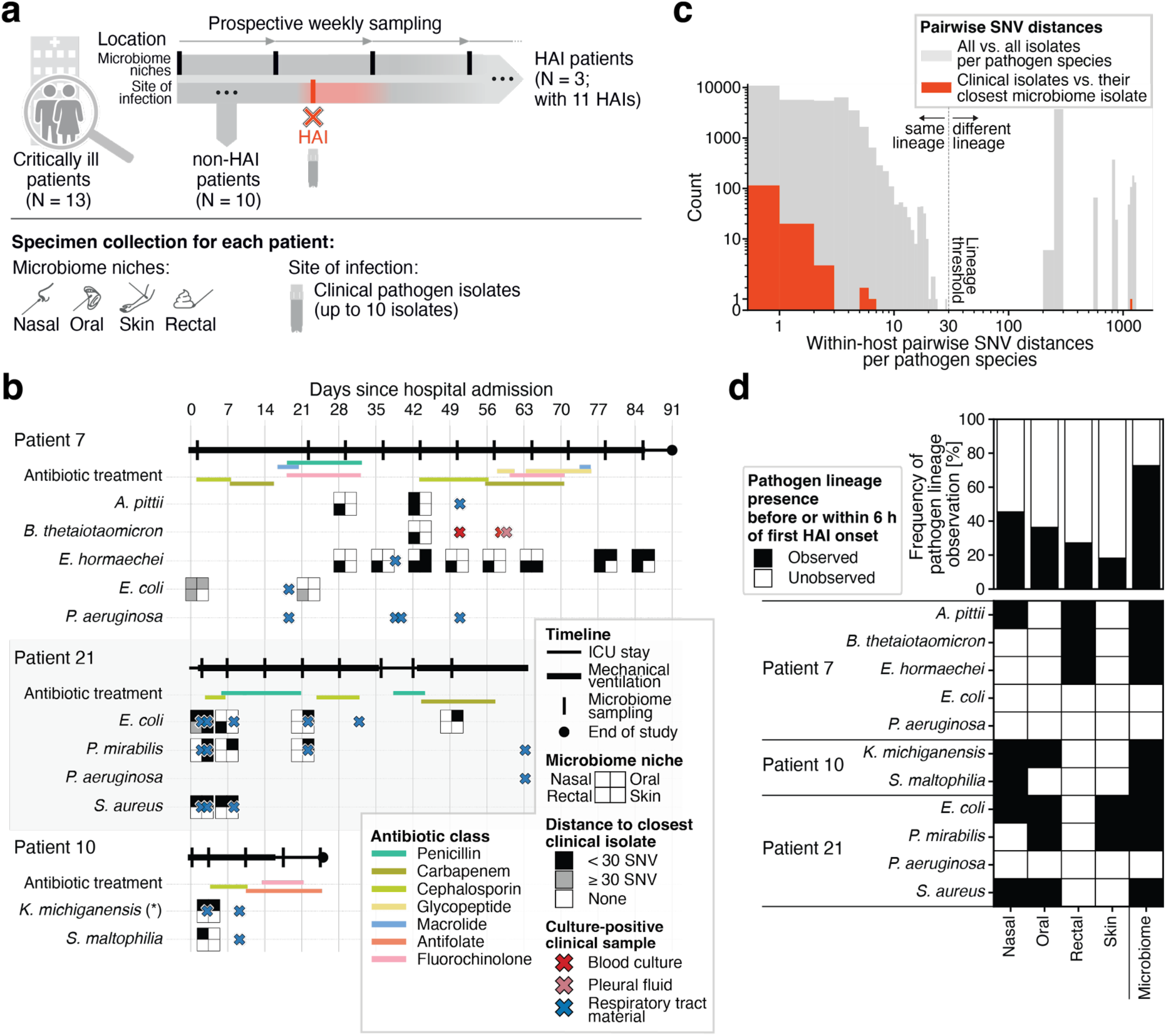
Prospective, longitudinal sampling of critically ill patients reveals body-wide lineage dissemination of opportunistic pathogens during healthcare-associated infection. **a.** Schematic representation of the prospective sampling established for this study. 13/25 (52%) eligible, critically ill patients were successfully enrolled for analysis (**Figure S1**). Prospective native microbiome samples (nasal and oral cavity, cubital fossa, and rectum) were collected weekly starting within 6-51 h after hospital admission (patient 7: 29 h; patient 10: 6 h; patient 21: 51 h). Upon an identified HAI up to 10 viable isolates of the pathogenic species were collected (see Methods). One or more HAI were detected in 3 of the 13 patients. **b.** Timelines of each HAI patient, which start at hospital admission and state periods of mechanical ventilation (thick horizontal line), timepoints of microbiome sampling (black vertical lines), and end of study (discharge from ICU or reaching the end of maximum enrollment period [90 days]; black dot). Colored horizontal lines show antimicrobial treatment, grouped by antibiotic classes. HAI onset timepoints defined as culture-positive clinical samples (blood culture [red], pleural fluid [lightred], and respiratory tract material [blue]) are shown as crosses. Squares are shown at every timepoint at which the respective pathogen species has been detected in at least one microbiome niche. Filled, black quadrants indicate that at least one isolate was observed to be closely related (less than 30 core-genome SNVs; see Figure 1c) to an isolate from the site of infection. If no such relationship was identified, the quadrants are filled in grey; if the species was not found in that microbiome niche, the quadrants are white. For patient 7, two consecutive microbiome sampling timepoints are missing (day 8 and 15). For patient 21, weekly microbiome sampling was discontinued after 49 days. A culture-positive gastric juice sample on day 3 in patient 10 for *K. michiganensis* (marked with an asterisk behind the species name) was detected coinciding with the microbiome sampling and infection identification. **c.** Core-genome pairwise SNV distances between all isolates per pathogen-patient pair. Comparison of isolates (grey) shows the majority of isolate pairs are less than 30 SNVs apart (indicated by the vertical dashed line), a degree of divergence plausible to accumulate during within-host colonization, which was therefore used to cluster isolates into lineages. Isolate pairs ≥ 30 core-genome SNVs apart were classified as separate lineages. Pairwise distances of every clinical isolate to their closest microbiome isolate (red) reveal that most of the clinical isolates belong to a lineage also present within the microbiome except for the *E. coli* ST95 in patient 7 (1,171 core-genome SNVs apart). Both axes are shown in log-scale and the bin size on the x-axis is one for < 100 SNV differences and 50 for ≥ 100 SNV differences to improve visibility on log-scale. **d.** Microbiome colonisation by the pathogen lineage before or within (≤ 6 h) HAI onset. Heatmap (bottom) shows the observation of the pathogen lineage (black) per species and microbiome niche as well as the collapsed, body-wide observation (microbiome) until 6 h after HAI onset. In the case of multiple pathogen lineages per pathogen-patient pair (*S. aureus* of patient 21), all pathogen lineages have been detected in at least one microbiome niche. The barplot on top shows the frequency of the identified microbiome colonization of the pathogen lineage across the eleven pathogen-patient pairs.

Culture-based isolate sequencing enables the identification of closely related isolates collected from the microbiome as well as the site of infection (**Figure S3**). We sequenced all clinical isolates collected from the site of infection and up to 20 isolates per pathogen species of each microbiome niche and timepoint, generating in total 755 genomes (**Figure S4**). First, comparing the isolates’ genomes collected from the site of infection shows high clonality within the reconstructed populations. We identified 0 to 12 pairwise core-genome single-nucleotide variant (SNV) differences along the core genome for all pathogens but *S. aureus.* Instead, *S. aureus* shows evidence for infection with two lineages separated by more than 200 pairwise core-genome SNVs (**Figure S5**). For all six HAI cases where the pathogen species was sampled repeatedly at the site of infection during hospitalization the observation of clonality holds up (**Figure S5**), suggesting persistence within the host. Second, extending the analyses to the genomes collected from the microbiome, in eight out of nine (89%) pathogen-patient pairs we identified a closely related genotype in the patients’ microbiome with 0 to 6 pairwise core-genome SNVs differences (**Figure 1c**). Subsequent phylogenetic reconstruction of all pathogen-patient pairs (**Figure S6 - S14**) confirms the close genetic relationship between isolates from the site of infection and the microbiome in those eight HAIs - in line with co-colonisation or translocation to-or-from the microbiome. Importantly, in all eight pathogen-patient pairs with microbiome culture success of the opportunistic pathogen lineage we identified the lineage prior to or within 6 h of HAI onset from a niche other than the site of infection (**Figure 1d**), pointing at a role of the microbiome during HAI.

Within the microbiome the majority of isolates showed 0-28 pairwise core-genome SNV differences in line with a single colonization event ^29,30^. Accordingly, we group closely-related isolates with 28 or less pairwise differences into distinct lineages (**Figure 1c**). We also found evidence for the co-colonization by different lineages of the same species within a patient. For *E. coli* (patient 7 and 21), *P. mirabilis* (patient 21), and *S. aureus* (patient 21) we collected isolates with more than 200 pairwise core-genome differences - too distant for a single colonization event (**Figure 1c, Figure S8, Figure S12 - S14**). For example, in patient 7, *E. coli* ST131, an extraintestinal pathogenic sequence type ^31^ was identified causing a ventilator-associated pneumonia with no representative cultured from the patient’s microbiome. Instead, sequence types 58, 95, 117 and 4260 co-colonized the patient’s gastrointestinal tract (**Figure S8**). Whereas in patient 21 *E. coli* sequence types 23 and 16602 have been cultured from the microbiome, but only the former has been linked to the patient’s hospital-associated pneumonia (**Figure S12**). Further, the *S. aureus* infection within patient 21 is caused by sequence types 45 and 97, and both are observed within the patients’ oral and nasal microbiome (**Figure S14**) consistent with the fact that 72-86% of *S. aureus* infections arise from the nasal microbiome ^32,33^. Interestingly, for 63% (5/8 HAI) of the pathogen lineages identifiable within the microbiome before or within 6 h of HAI onset, the lineage has been cultured successfully in multiple microbiome niches (**Figure 1d**). Such a pattern is in agreement with either a recent colonization of the patient’s body from the clinical environment or reflects a momentary footprint during bacterial translocation within the patient’s body raising the question about the timing of colonization.

### Microbiome colonization by a subset of pathogen lineages precede HAI onset or even hospitalization

The reconstructed diversity of each pathogen lineage within the microbiome may allow the inference of the colonization timing, and to estimate whether or not the pathogen was acquired before or after hospital admission. Here we infer for two pathogen lineages that colonization precedes HAI detection, of which one lineage has an inferred colonization time before hospitalization. In a first step, we leveraged the observed mutations over time to estimate the molecular clock for each pathogen lineage. However, we only identified a clock-like root-to-tip signal for *E. hormaechei* in patient 7 (2.765 SNVs/Mb/year [95% CI: 2.22-3.31]; **Figure S15**). For all remaining pathogen species we instead considered using published molecular clocks. To validate the applicability of published molecular clocks, we compared the observed mutational spectrum of the pathogen lineages with the mutational spectrum of published genomes. This revealed for *E. coli* ST23 (patient 21) a significant deviation in the mutational spectrum, which was subsequently omitted from analysis (**Figure S16**). Of the remaining five species (*A. pittii, B. thetaiotaomicron, K. michiganensis, P. mirabilis, S. aureus*), we only identified molecular clocks for *S. aureus*. Importantly, mutation rates of a species can vary, depending for example on the population density, environmental stress (e.g. antibiotics) or genetic background ^34–36^. To minimize confounding effects in the utilized rate, we calculated the molecular clock of *S. aureus* using published longitudinal isolate sequencing data part of the same sequence type – ST97 (2.548 SNVs/Mb/year [95% CI: 2.241, 2.856, r^2^ = 0.574]; **Figure S17a**).

For both, the *E. hormaechei* and *S. aureus* ST97 pathogen lineages, we inferred the time to the most recent common ancestor (TMRCA) based on the ascertained diversity within the microbiome at the earliest time point with sufficient data (≥ 8 isolates), which was either before HAI (*E. hormaechei*: 57 h before HAI onset) or within 6 h of HAI onset (*S. aureus* ST97: 2 h after HAI onset). For *S. aureus* ST97, the TMRCA inference estimates colonization and diversification within the microbiome of patient 21 already before entering the hospital (**Figure 2a**). Comparable results are obtained using an array of additional, published molecular clocks of *S. aureus* from various genetic and environmental backgrounds, corroborating our inference of pre-hospitalization colonisation (**Figure S17b**). In contrast, the *E. hormaechei* pathogen lineage shows evidence for acquisition prior to infection onset but after hospital admission (**Figure 2a**). Notably, incomplete reconstruction of the population diversity during culturing or recent population bottlenecks introduced for example by antibiotic treatment may bias the inference towards more recent events. A collector’s curve analysis suggests sufficient sampling of the timepoint used for inference, with the distance to the most recent common ancestor reaching a plateau for *S. aureus* and *E. hormaechei* (**Figure S18**). Nevertheless, we do not sample every low-frequency genotype (**Figure S19**); those, however, do not affect the TMRCA analysis. Altogether, we provide evidence for both HAI pathogen lineages colonizing the patients’ microbiomes preceding detection of infection, and *S. aureus* ST97 being present prior to hospitalization, underlying the importance of the human microbiome as a reservoir and translocation compartment for HAI pathogens within the two patients analysed.

**Figure 2:**
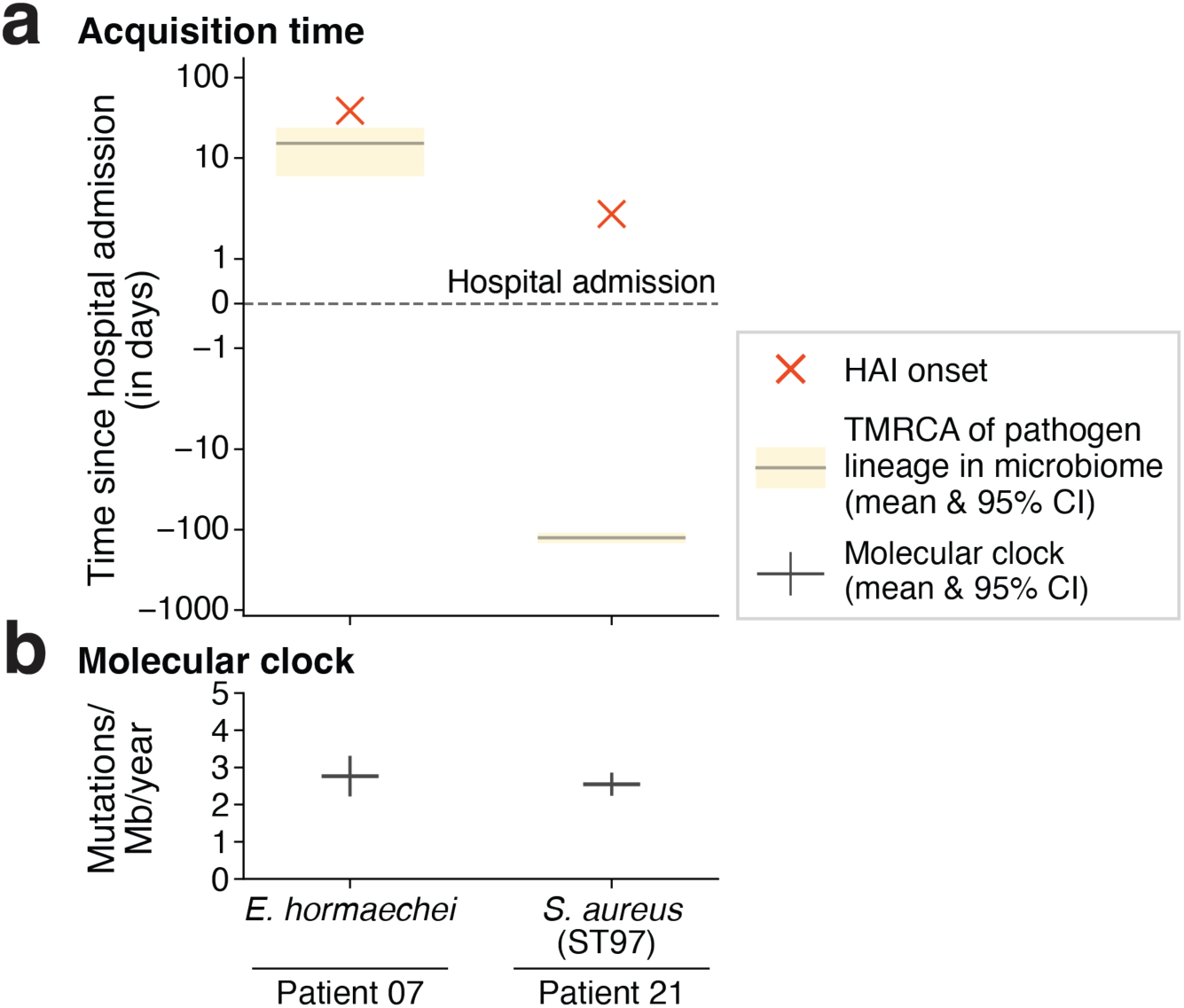
Estimating the time to the most recent common ancestor within the patient’s microbiome for two pathogen lineages. **a.** Inferred time to the most recent single-celled ancestor of the pathogen lineage (mean in grey; 95% confidence interval in yellow) in respect to clinical diagnosis (red cross). Time since hospital admission shown with a dashed line. Times were inferred using the earliest timepoint where eight or more isolates have been collected from the microbiome to approximate diversity, which was either before or within 6 h of HAI onset. Confidence intervals were calculated using 10,000 bootstraps of the observed genotypes combined with random sampling from a gamma distribution fitted to the molecular clocks. **b.** Molecular clocks and their 95% confidence interval used for the TMRCA analysis were inferred either from the patient data directly (*E. hormaechei* [**Figure S15**]) or from longitudinally collected *S. aureus* isolate-genomes part of the same sequence type infecting patient 21 (ST97) ^11^.

### Short-lived mutation in *E. hormaechei* associates with infection

Human commensal bacteria likely experience challenges for their survival at the site of infection triggered by the host’s immunity, treatment, and alterations in the environment. However, we only have a limited understanding about whether or not *de novo* mutations contribute to infection initiation, because prospective, longitudinal microbiome data collected before infection diagnosis is uncommon. Here, we leverage the reconstructed within-patient diversity of each pathogen lineage to investigate SNVs, insertions and deletions (indels), and mobile genetic elements (MGEs) associated with infection. We identified a mutation in *Enterobacter hormaechei*, of patient 7 that became associated with infection.

Patient 7 sequentially developed symptoms of ventilator-associated pneumonia (VAP) with purulent respiratory conversion and radiographic evidence of pulmonary infiltrates 39 days after hospitalization, followed by fever and leucocytosis. At the time of radiographic evidence, tracheal aspirate was collected and confirmed to be culture-positive for *Enterobacter hormaechei* and *Pseudomonas aeruginosa* at similar CFUs (**Figure 1b, Table S2**). While we were unable to culture *P. aeruginosa* from any microbiome sample, we successfully cultured 192 *E. hormaechei* isolates from the four sampled microbiome niches of patient 7 over a period of nine weeks (**Figure 1b**; **Figure 3a**).

**Figure 3:**
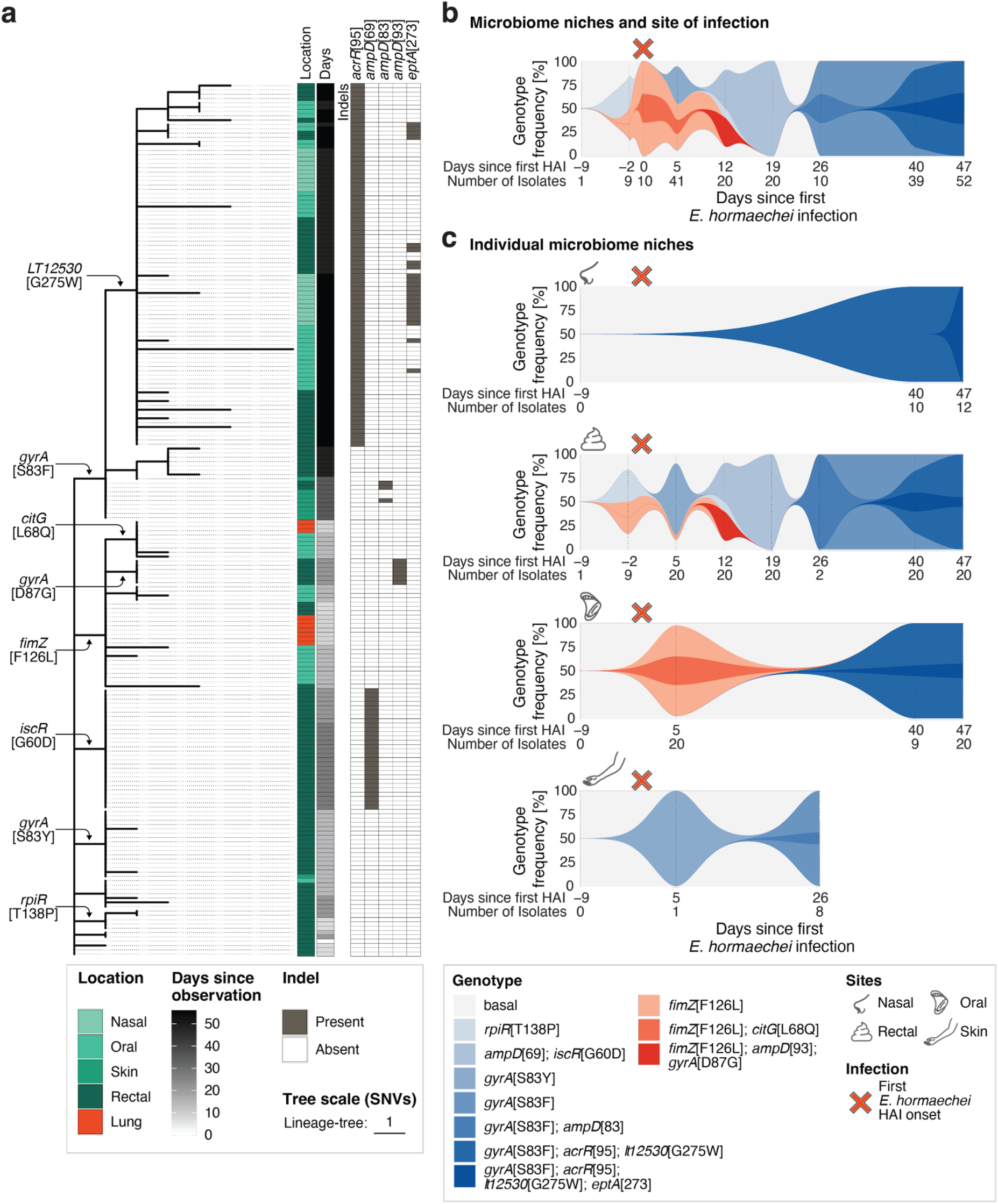
Short-lived *de novo* mutation *fimZ* [F126L] first detected in the gut becomes associated with infection but subsequently is replaced by independently, emerging genotypes linked to treatment resistance. **a.** Maximum-likelihood phylogeny of the 202 *E. hormaechei* isolates from patient 7 using the lineage-specific alignment. The tree, based on 61 SNVs, is rooted on GCF_003408555.1 and the tree scale bar represents one SNV. The microbiome isolation source is shown in shades of green or red for isolates collected from the site of infection. Timepoints since first isolation (in days) are shown in grey scale. SNVs with a frequency shift ≥ 30% between observations are annotated on the respective branches. Indels with a genotype frequency shift ≥ 30% between observations are shown in a heatmap in grey if present in the respective isolate with the gene name and position of the mutated amino acid in squared brackets on top. **b-c.** Muller plots show *E. hormaechei* SNV- and indel-based genotypes with frequency shifts ≥ 30% for (**b**) body-wide (microbiomes and site of infection collapsed) and (**c)** niche-specific (nasal [top panel], oral [second panel], rectal [third panel], and skin [lower panel]) from first culture-success (−9 days prior infection detection) until patient 7’s last sampled timepoint (47 days). Frequencies were interpolated in log-space to model exponential growth, restricting to timepoints with at least one observed isolate. Each color represents a different genotype (shades of red indicate genotypes carrying the *fimZ* [F126L] mutation) and the sampled *E. hormaechei* infection timepoint is highlighted with a red cross above each plot.

Tracking the spatio-temporal diversity of *E. hormaechei* across patient 7 identifies a short-lived mutation associated with infection. The earliest *E. hormaechei* isolate had been collected from the patient’s gut microbiome nine days prior to VAP onset. Next, two days before VAP detection we identified isolates in the gut microbiome carrying a non-synonymous *de novo* mutation (F126L) in the type I fimbriae regulator gene *fimZ* ^37^, which became the exclusive genotype associated with infection. Five days later, *E. hormaechei* became detectable on the skin, oral cavity and gut, with the F126L genotype culturable in the latter two niches and at 12 days post-infection the F126L genotype was cultured for the last time - from the gut microbiome (**Figure 3a**, **Figure 3c**). At the same time, independent, closely-related genotypes carrying the wildtype allele - the ancestral F126 - were observed within different niches carrying SNVs and indels in known resistance-conferring genes, for example *ampD* ^38^ and *gyrA* ^39^, associated with multiple bodywide sweeps (**Figure 3**). Strikingly, both genes showed strong signatures for parallel evolution with eleven independent mutations within *ampD* (p-value = 6.7*10^-38^, Poisson test) and five independent mutations within *gyrA* (p-value = 3.3*10^-10^, Poisson test), possibly elicited by the treatment regime of the patient (see below). Notably, tracheal aspirates of patient 7 were culture-positive for *E. hormaechei* three and 39 days later together with clinical parameters in line with VAPs (**Figure S2,** Methods), but no isolates were collected leaving their precise genotype obscure. No *fimZ* [F126L] carrying genotype was isolated at the latter timepoint (39 days after initial infection) from the microbiome, leaving infection with any of the newly arisen genotypes linked to treatment resistance plausible, given their widespread prevalence across the body at the time. In addition, tracheal aspirates were culture-positive for *P. aeruginosa* at another four timepoints, some of those alongside *E. hormaechei*, suggesting a polymicrobial infection driving the clinical VAP manifestation (**Figure S2**, **Table S2,** Methods). While the contribution of the different pathogens to the VAP cannot be discerned and despite the subsequent *E. hormaechei* infecting genotypes undefined, the identified association of the *fimZ* [F126L] mutation with the initial *E. hormaechei* infection may point to a possible (patho-)adaptive advantage.

### HAI-associated *E. hormaechei fimZ* [F126L] affects biofilm formation, adhesion and virulence

Type I fimbriae are short pili used by *Enterobacteriaceae* to attach to surfaces or cells; they are important for biofilm formation and adhesion, both of which are linked with virulence, for example in uropathogenic *E. coli* causing UTIs ^40–43^. FimZ activates type I fimbriae expression by binding to the *fimA* promoter of the operon FimAICDHF (**Figure 4a**) leading to the biosynthesis of fimbriae, helical structures of FimA monomers on the outer membrane with the adhesin FimH at its tip ^44^. Given that the observed amino acid changing mutation F126L is located at the protein surface within a negative patch (**Figure 4b**) we hypothesized that this mutation may alter fimbriae expression and *E. hormaechei*’s ability to colonize the lung.

**Figure 4:**
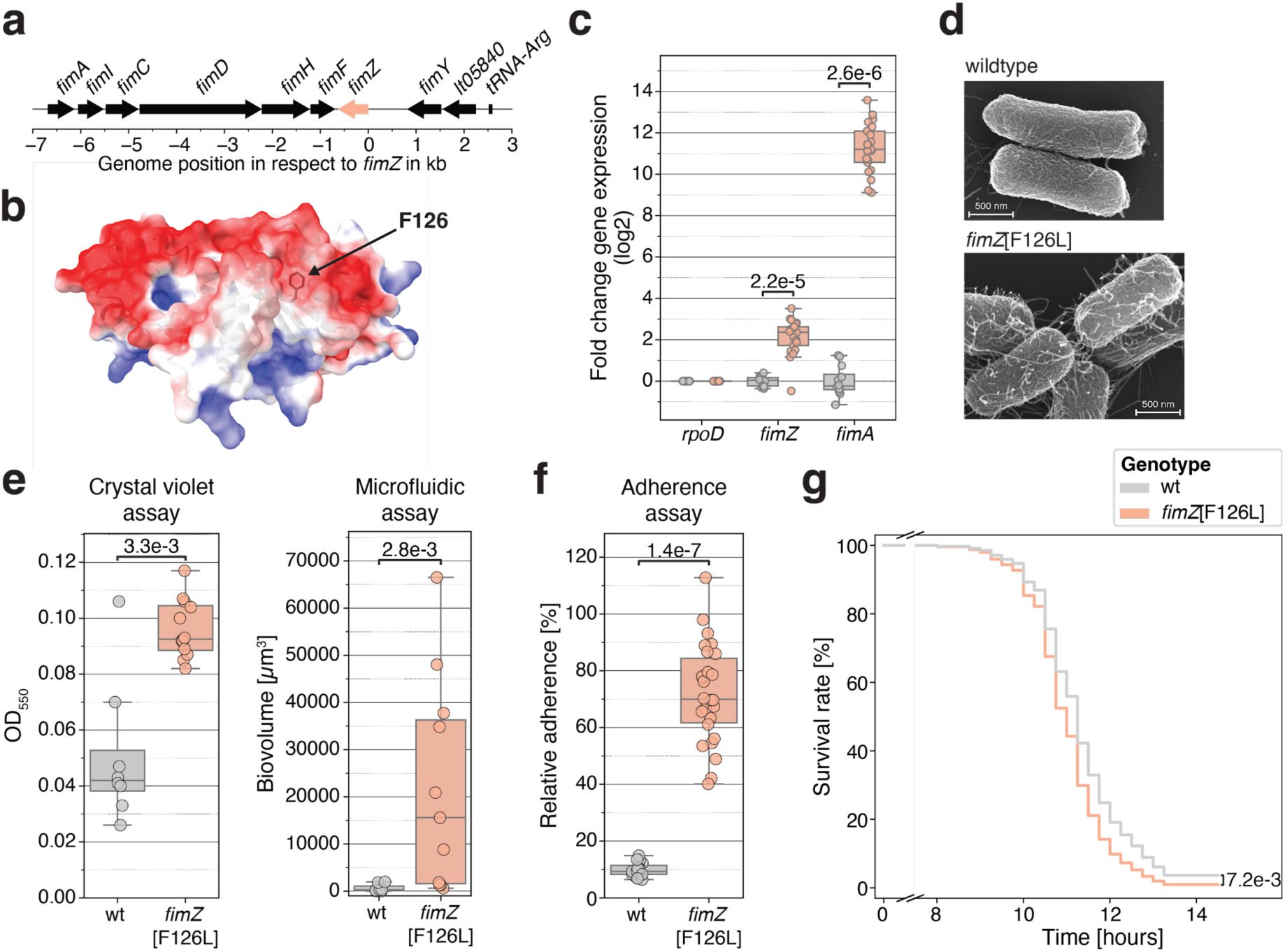
Infection-associated *E. hormaechei fimZ* [F126L] shows elevated fimbrial expression, biofilm formation, adhesion, and virulence. **a.** Genetic orientation (arrows) of the genes within the type I fimbrial operon observed in the patient-specific *E. hormaechei* genome. The *fimZ* gene is shown in salmon and the other genes are shown in black. Each gene is annotated above. **b.** Predicted 3D structure of FimZ carrying the wildtype allele. The surface is color-coded by its electrostatic charge (negative [red], positive [blue]). Location of the amino acid F126 is indicated by an arrow. **c.** Transcriptional expression of the housekeeping gene *rpoD* (RNA polymerase sigma factor RpoD), the transcriptional regulator *fimZ*, and the structural fimbriae gene *fimA* (type 1 fimbrial major subunit FimA). Genotypes carrying the wildtype, ancestral *fimZ* [F126] (P07_2048, P07_2055), or genotypes with the *fimZ* F126L] mutation from the microbiome (P07_2052, P07_2237) or the site of infection (L00071_01_1, L00071_02_1) were used. Presented results are collapsed across two growth phases (early- and late-log) each measured in three biological replicates per isolate. Statistical differences were assessed using a Mann-Whitney U test. **d.** Scanning electron microscopy images of the ancestral, wildtype (P07_2048) and derived *fimZ* [F126L] genotype (L00071_01_1) from late-log phase. Scale-bar represents 500 nm. **e.** Biofilm formation differences between wildtype *fimZ* [F126] genotype and *fimZ* [F126L] mutant genotype isolates (same as in **c**) at stationary conditions (Crystal violet assay; left plot) and under constant flow in minimal media (Microfluidic assay; right plot). Statistical analysis was done as in (**c**). **f.** Adherence of isolates with the wildtype or *fimZ* [F126L] genotype (same as in **c**) to human lung epithelial cells (Calu-3) *in vitro*. Adherence capacity was normalized to the inoculum. Statistical analysis was done as in (**c**). **g.** *Drosophila melanogaster* survival assay of male *Relish^E^*^20^ iso flies systemically infected with either a wildtype genotype (P07_2048, P07_2055) or a *fimZ* [F126L] mutant genotype (P07_2052, L00071_01_1). Replicates with significantly different initial bacterial inoculation levels were removed (based on CFUs; **Figure S23a**). The survival curves were fitted per genotype (*fimZ* [F126L]: 118 flies; wildtype: 159 flies) using the Cox proportional-hazards model and significance was assessed via the integrated Wald test including the experimental replicates as covariate. The x-axis is broken (0.5-7.5 h) to remove the inoculation period. Individual Kaplan-Meier curves are shown in **Figure S23b**.

Here, we deploy *in vitro* and *in vivo* assays to test expression, biofilm formation, adherence, and virulence utilizing isolates directly collected from patient 7 that carry the *fimZ* [F126L] mutation and compare them to otherwise isogenic or closely related isolates carrying the ancestral allele *fimZ* [F126], defined as wildtype isolates hereafter (**Figure S20**). First, comparing the gene expression of *fimZ* itself as well as of *fimA*, the major structural gene of type I fimbriae, shows both genes are consistently elevated in the isolates carrying *fimZ* [F126L] compared to wildtype isolates carrying the ancestral allele (p-value [*fimZ*] = 2.2*10^-5^; p-value [*fimA*] = 2.6*10^-6^, Mann-Whitney U test; **Figure 4c**), an effect independent of growth phase. Notably, the higher expression of *fimZ* and *fimA* translates into a distinct phenotype, where the *fimZ* [F126L] mutant shows a hyper-piliated phenotype with more fimbriae-like structures on the cell surface compared to wildtype isolates (**Figure 4d**). Second, biofilm formation was measured *in vitro* using a crystal violet and a microfluidic assay identifying cell aggregation to an abiotic surface under static and media flow conditions, respectively ^45–47^. Both complementary assays showed a consistent increase in cell aggregation for different isolates carrying the *fimZ* [F126L] mutation versus different isogenic wildtype isolates (p-value [crystal violet assay] = 3.3*10^-3^ ; p-value [microfluidic assay] = 2.8*10^-3^, Mann-Whitney U test; **Figure 4e**). Third, adhesion to human lung epithelial cells (Calu-3) was quantified *in vitro*, demonstrating the *fimZ* [F126L] isolates exhibit increased adherence compared to wildtype isolates (p-value = 1.4*10^-7^, Mann-Whitney U test; **Figure 4f**). Notably, the growth rates between the *fimZ* [F126L] mutant and the wildtype are similar and do not contribute to the observed differences providing no evidence for a fitness cost effect by *fimZ* [F126L] (**Figure S21**). Altogether, those results show that the *fimZ* [F126L] mutation elevates *in vitro* fimbriae production increasing biofilm formation and adhesion to the human lung epithelium.

Encouraged by the observed phenotypic differences *in vitro*, we tested whether the hyper-piliated *fimZ* [F126L] genotype also translates into increased virulence *in vivo* compared to the wildtype isolates harboring the ancestral allele. Therefore, we used a systemic infection assay of wildtype and immunocompromised (*Relish^E^*^20^ iso) *Drosophila melanogaster* ^48^. *Relish^E^*^20^ iso flies have a deficiency in the humoral Imd pathway important for an antimicrobial response, rendering them sensitive to gram-negative infections. While wildtype flies exhibited no survival defects between tested genotypes (p-value = 0.62, Wald test; **Figure S22**), we observed a moderate but significantly increased disease progression of immunocompromised *Relish^E^*^20^ iso flies infected with *fimZ* [F126L] isolates compared to wildtype isolates (p-value = 7.2*10^-3^, Wald test; **Figure 4g**). The results indicate increased virulence of the hyper-piliated *fimZ* [F126L] mutant in a gram-negative-sensitive fly infection model. In summary, we show a hyper-piliated *E. hormaechei* phenotype linked to functional differences attributable to the short-lived, *de novo* mutation F126L in *fimZ*, which may possibly point to an adaptive pathway of the pathobiont during infection.

### Global clinical *E. hormaechei* analysis suggests *fimZ* [F126L] is rare

While the phenotypic differences measured *in vitro* and *in vivo* suggest an adaptive advantage for the *E. hormaechei fimZ* [F126L] mutant to establish a respiratory tract infection in patient 7, it remains unknown whether it represents a common adaptive path for infections within the respiratory tract among hospitalized patients. Due to the limited sample size of our dataset in which we only observed a single patient infected with *E. hormaechei*, precluding any repeated observation necessary to deduce whether mutations in *fimZ,* and especially F126L, are common or rare during infection. To evaluate the frequency of mutations in *fimZ* within the clinical context, we leveraged 480 publicly available genomes of *E. hormaechei* isolated from human respiratory samples collected across 23 countries spanning all continents except Antarctica (**Table S3**). We identified a total of 79,610 SNVs within these genomes without a phylogeographic structure (**Figure S24a, Data S1**). We observe four non-synonymous mutations in *fimZ* (E11Q, G36A, V155A, A171V) among five (1.04%) analysed genomes, whereas seven synonymous mutations were identified. Using the dN/dS ratio, a canonical test for natural selection, suggests that *fimZ* evolves under purifying selection (dN/dS*_fimZ_* = 0.252) residing in the central range of the genome-wide distribution (83rd percentile; **Figure S24b**). Noteworthy, dN/dS measurements can be biased when sampled within a population ^49,50^. However, the utilized collection of divergent lineages of *E. hormaechei* (median(DMRCA) = 4644.5 SNVs) combined with the strong departure from the expected rate of the *E. hormaechei* lineage of 2.5:1 non-synonymous to synonymous mutations, suggests that *fimZ* evolves under evolutionary constraint and does not usually carry amino acid-changing mutations associated with respiratory tract infections. This suggests the observation of the *E. hormaechei fimZ* [F126L] mutant in patient 7 is rare despite its phenotypic implication.

### Multiple, body-wide sweeps of treatment-resistance conferring mutations lead to loss of *fimZ* [F126L]

The *fimZ* [F126L] mutation becomes undetectable as independently emerging mutations in genes facilitating antimicrobial resistance become dominant. To explore the antibiotic susceptibility landscape over time, we combine the known treatment history with antimicrobial susceptibility tests using the collected *E. hormaechei* isolates to identify whether or not the repeated, independent sweeps of novel *E. hormaechei* genotypes on patient 7 are linked to resistance (**Figure 3**). Importantly, the initially detected *E. hormaechei* genotype in the gut as well as the emerging, closely-related *fimZ* [F126L] mutant are susceptible to the antibiotics administered to patient 7, suggesting the association of F126L with infection is not treatment induced (**Figure 5**). However, within 12 days after detection of *E. hormaechei* in VAP and subsequent ceftazidime treatment (3^rd^ gen. cephalosporine beta-lactam antibiotic) several genotypes emerged. Among these we observed various SNVs in *gyrA* as well as gene-truncating indels in *ampD*, some of which are linked to *fimZ* [F126L] (**Figure S9**). The tested isolates with *ampD* truncations (position 68, 93, or 115) mediate increased resistance against piperacillin/tazobactam and cefotaxime by more than 42-fold, exceeding EUCAST clinical breakpoints, and the isolates carrying *gyrA* SNVs show a more than 5.9-fold observed MIC increase against fluoroquinolones (**Figure 5**). Eleven days after the *E. hormaechei* VAP an independent genotype of *E. hormaechei* not carrying *fimZ* [F126L] but an *iscR*[G60D]-*ampD-*truncation emerges, which mediates beta-lactam resistance against cefotaxime and piperacillin/tazobactam (**Figure 5**), and with increased MIC against Meropenem (**Figure S25**) following treatment with the beta-lactam antibiotic ceftazidime. At day 18 after VAP solely the *iscR*[G60D]-*ampD-*truncation genotype is detectable suggesting a treatment-associated selection, concurrent with *fimZ* [F126L] becoming subsequently undetected (**Figure 3a**). During the following week, the patient was treated with vancomycin, imipenem and ciprofloxacin, coinciding with the establishment of a novel, independent genotype carrying a non-synonymous mutation in *gyrA* [S83F] associated with ciprofloxacin resistance (**Figure 3a**, **Figure 5, Figure S9**). Following a culture-negative time point, we observed 40 days post-infection two different genotypes carrying *gyrA* [S83F]; one carried an additional SNV in the promoter region of *romA* and *acrR,* and another with a gene-truncating indel in the coding sequence of *acrR* (amino acid location 94), the regulator of the multidrug efflux pump AcrAB-TolC ^51^, both associated with increased ciprofloxacin resistance of 2- to 3.9-fold (**Figure 5**). This occurrence also coincided with another unsampled episode of VAP caused by *E. hormaechei* in patient 7 (see Methods, **Figure S2**). A week later, at the last sampled time point, the *gyrA*[S83F]-*acrR*[94]-truncation was observed within all identified isolates and sampled from all tested microbiome niches but skin (**Figure 3c, Figure S9**). Five days later, patient 7 ultimately reached the maximum enrollment of 90 days, presumably colonized with an *E. hormaechei* population lacking the *fimZ* [F126L] genotype but being highly resistant to ciprofloxacin.

**Figure 5:**
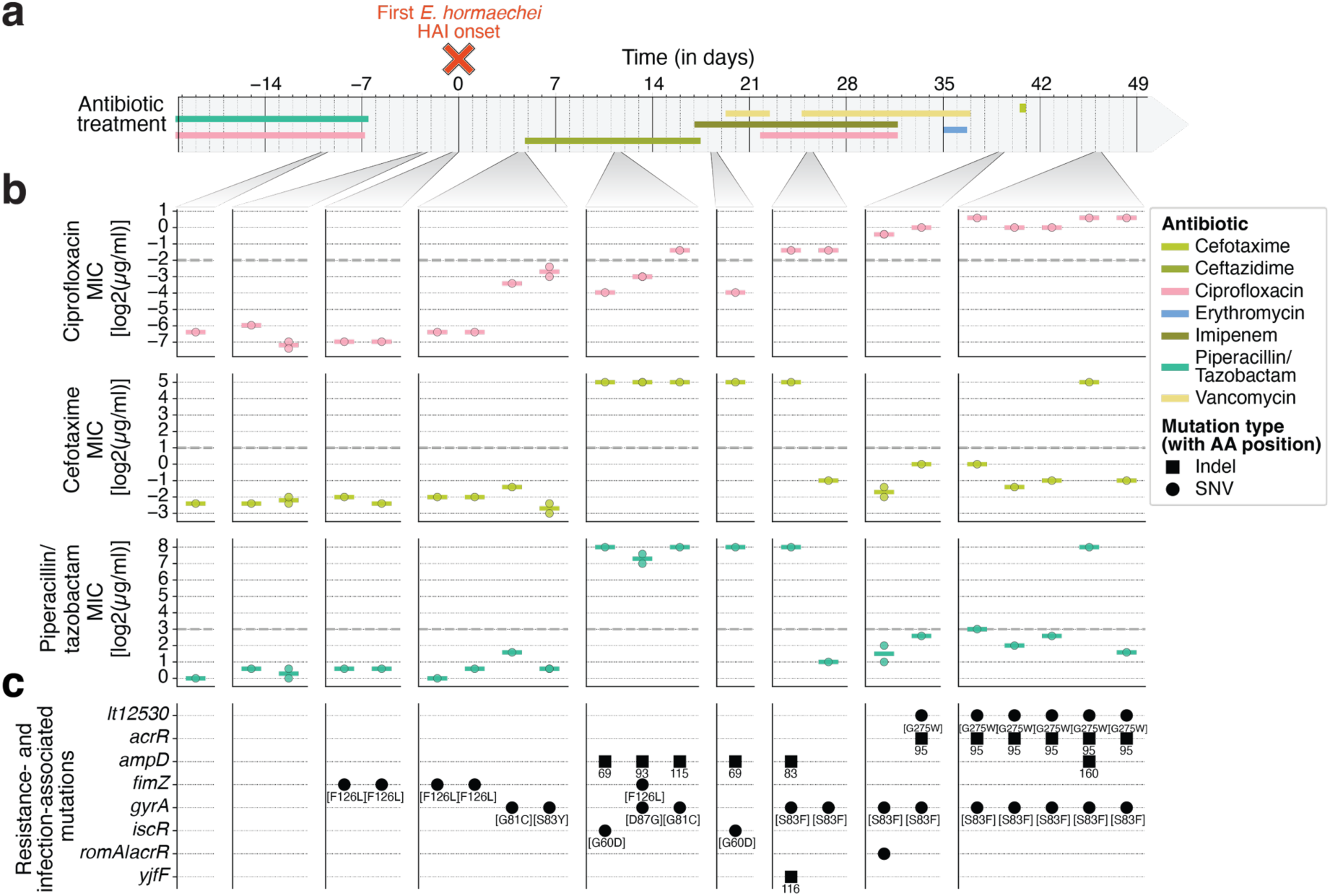
Within-patient adaptation to antibiotic treatment leads to removal of the *fimZ* [F126L] mutant and to the establishment of a ciprofloxacin-resistant *E. hormaechei* population within patient 7. **a.** Antibiotic treatment episodes of patient 7 shown, beginning 20 days prior to the first infection with *E. hormaechei* (red cross) until 3 days after the last microbiome sample of patient 7. **b.** Minimum-inhibitory concentrations (MIC) of selected isolates representing major branches (**Figure S9**) against antibiotics used for treatment. Resistance was defined following the EUCAST Clinical Breakpoints Tables (v15.0; grey dashed lines; see Methods) and only antibiotics for which resistance breakpoints were exceeded are shown: fluoroquinolone (ciprofloxacin), and two beta-lactam (cefotaxime; piperacillin/tazobactam); see **Figure S25** for all tested antibiotics. Sampled timepoints are grouped and columns represent individual genotypes (summarized in [**c**]) of which one or two isolates were tested. Each point represents one isolate (up to two isolates per genotype) and the horizontal line represents the mean of the MIC. **c.** Resistance- and infection-associated SNVs and indels for each genotype are shown. Other variants are removed for clarity (see **Figure S25** for all SNVs and indels). Below mutations the amino acid change is stated for SNVs and the amino acid location for the start of indels. One intergenic mutation, a SNV, located between divergently transcribed genes (*romA*|*acrR*) was located 68 nt downstream of *romA* and 124 nt upstream of *acrR*.

## Discussion

Here we mapped the evolutionary history of opportunistic pathogens causing acute infections in three critically ill patients using native, prospective, longitudinal sampling coupled with whole-genome sequencing. For the majority of infections, the clonal or near clonal relationship among collected clinical isolates showed infections derived from a single genotype. Moreover, we identified closely related isolates in 73% of the infections within the patients’ microbiomes, either before or at the onset of HAI symptoms, emphasizing that co-colonization of the patient’s microbiome accompanies infection. Leveraging the reconstructed population diversity from microbiomes for molecular dating implies, for an *S. aureus* infection, that acquisition occurred prior to hospitalization. This highlights the role of the patient’s microbiome as a potential reservoir for opportunistic pathogens consistent with previous studies ^19,33,52^ Moreover, TMRCA inference is dependent upon the molecular clock, which can vary between lineages of a species and depends upon environmental pressures (for example host immunity or population density). Hence, we solely perform TMRCA inference for pathogen lineages where we either derive the molecular clock directly from our patient data (*E. hormaechei*) or from longitudinally collected isolates of closely related strains part of the same sequence type (*S. aureus*). Any inferred TMRCA, however, only provides lower bound estimates for the colonization timing of the microbiome, because incomplete niche sampling, culture biases and antimicrobial treatments interfere with the reconstruction of the complete lineage diversity.

Disentangling evolutionary trajectories revealed a *de novo* mutation in the type I fimbriae regulator gene *fimZ* among *E. hormaechei* isolates in one patient, first detected within the gut microbiome that became associated with pneumonia. The mutant showed increased biofilm formation and lung epithelia adhesion *in vitro* and elevated virulence *in vivo*, suggesting a pathoadaptive role. While our observation is restricted to a single patient, the analysis of 480 public isolates from human respiratory samples shows the genetic stability of FimZ at the amino acid level, including the absence of the *fimZ* [F126L] mutation, rendering *fimZ* [F126L] rare during colonization and infection of the respiratory tract. Interestingly, Frutos-Grilo et *al.* (2023) has identified type I fimbriae-mediated adherence of *Enterobacter cloacae* to bladder, kidney and lung epithelial cells *in vitro* with reduced adherence to epidermal and colon cells ^53^. This supports a niche-specific colonization advantage of the *fimZ* [F126L] strain within the lung but little effect within the intestine, its commensal environment. Despite type I fimbriae being widespread across *Enterobacteriaceae* and recognized as a crucial virulence factor for uropathogenic *E. coli* and *Klebsiella pneumoniae* ^40,41,54^, it has been demonstrated that they are regulated differently and have different specificities ^54,55^. This highlights the difficulty in identifying common pathoadaptive strategies in orthologous genes across different opportunistic pathogenic species ^9^. Ultimately, the *fimZ* [F126L] strain is either cleared through antibiotic therapy or replaced by independent *de novo* genotypes shown to mediate elevated antimicrobial drug resistance which are sweeping consecutively through the *E. hormaechei* population and across the patient’s body. Moreover, alongside *E. hormaechei*, *P. aeruginosa* was detected within the patient’s tracheal aspirate and it remains unknown whether both or one of the two were driving the infection. However, the combination of genetic, experimental, and spatio-temporal evidence corroborates a pathoadaptive role for the *fimZ* [F126L] *E. hormaechei* strain in this patient during acute infection that is, both, rare and short-lived. This exemplifies the challenges to identify pathogen adaptations during acute infection when they follow along independent evolutionary trajectories rather than common and conserved genetic mechanisms.

While the mechanisms behind HAI are multifactorial, including invasive devices, underlying disease severity, or comorbidities, our findings support the role of the human microbiome during acute infections and support the hypothesis that short-sighted evolution gave rise to an adapt-and-die mutation associated with disease - in a single patient ^28,56^. While the logistical challenges of prospective clinical sampling and culture-based single-isolate sequencing pose a throughput bottleneck resulting in a small HAI cohort and the potential undersampling of minority genotypes and lineages, our proof-of-principle study proves successful in identifying a transitional genetic association within the critical window leading up to infection. Hence, it provides advantages over metagenomics or genome-wide association studies using single, clinical isolates ^21,57^. Nevertheless, additional efforts including larger patient cohorts are necessary to further disentangle adaptive within-host evolution linked to disease to determine the contribution of genetic variants on acute infections.

## Methods

### Study cohort

The target cohort of this study were adult critically ill patients, classified as patients who are at least 18 years of age with an emerging organ dysfunction, indicated by the increase of at least two points in the Sequential Organ Failure Assessment (SOFA) score, and an expected ICU stay of at least 72 h. Patients were excluded who have been hospitalized within the last 30 days before the current clinical admission, received antibiotic treatment within the last 30 days or have been under current antibiotic treatment for more than 24 h, as well as pregnant or breast-feeding women. A total of 174 critically ill patients admitted to the interdisciplinary medical intensive care unit (ICU) at the University hospital Leipzig (UHL) between June 2021 and September 2021 were assessed by the clinical staff for eligibility according to a study protocol approved by the ethics commission board of the University of Leipzig (DRKS00024777). Informed consent was obtained by the patient or their legally authorized representative, and 14 suitable patients were successfully enrolled into the study (**Figure S1**). Clinical data of each patient was collected pseudonymized in a database ^58^ throughout their hospital stay. Those data included the patient’s age, sex, underlying diseases and comorbidities, organ support therapy, clinical scores, treatment, and suspected and diagnosed infections with their clinical laboratory results. Metadata for each patient is presented in **Table S1**.

### Microbiome sampling

During the ICU stay, the microbiome of each enrolled patient was sampled weekly as part of the clinical routine screening for multiresistant pathogens. Prospective microbiome sampling for the three HAI patients was initiated 6-51 h after hospital admission: Patient 7 (29 h after hospitalization), patient 10 (6 h after hospitalization), and patient 21 (51 h after hospitalization). The delay in patient 21 occurred because the patient was first hospitalized for 46 h at a ward not participating in the study before being transferred to the ICU. Four different microbiome niches were sampled using sterile flock swabs: the nasal and oral cavity, rectum, and skin (cubital fossa) alongside a negative control (air-swab). Before sampling of the skin, the swab was moistened with sterile PBS. After sampling, each swab was transferred into 1.8 ml cryotubes containing 1 ml 30:70 glycerol:PBS, vortexed for 30 s and stored at - 80°C within 4 h after sampling to preserve sample viability. If a patient required mechanical ventilation and had a feeding tube inserted, a sample of the gastric aspirate was taken and stored unaltered in a 1.8 ml cryotube at -80°C.

### Definition of healthcare-associated infection

Healthcare-associated infections (HAIs) were generally defined as diagnosed infections occurring at least 48 hours after hospital admission, based on the American Thoracic Society (ATS) definition of hospital-associated pneumonia (HAP) ^59^ and the Linder-Mellhammar criteria of infection in sepsis for non-respiratory HAIs ^60^. The study focused on bloodstream infections (n = 1), pneumonia (n = 10), and urinary tract infections (none).

Hospital-associated pneumonia required at least two of the following criteria: a) abnormal temperature (< 36°C or > 38°C), b) leukocyte counts (< 4.0 Gpt/l or > 10.0 Gpt/l), and c) new purulent respiratory conversion and additional new radiographic evidence of pulmonary infiltrates. Ventilator-associated pneumonia was defined as hospital-associated pneumonia occurring after at least 72 h of invasive mechanical ventilation.

Healthcare-associated bloodstream infection required at least one culture-positive blood sample (two separate in case of a possible skin contaminant).

### Sampling of suspected healthcare-associated infections

Upon clinical suspicion of a HAI, in rare cases preceding the timepoint when all HAI criteria were met, a microbiological sample, marked as study material, was sent for culturing to the medical microbiology unit at UHL (see “Culturing and isolation of single pathogen colonies”). In case this provided culture-based evidence for an infection, the timepoint was defined as HAI onset. In case a pathogen was observed to be associated with a community-acquired infection (< 48 h post hospitalization) within the same patient, the pathogen was excluded in this patient from the analysis. Due to logistical challenges of the study corroborated by staff limitations during the COVID-19 pandemic, sampling was partially incomplete which led to the exclusion of patient 18 due to failure to retain the only clinical sample. It also caused the discontinuation of patient 21 after 63 days and resulted in an overall availability of 61.5% for isolates from diagnostic samples (**Figure S2, Table S1, Table S2**).

### Culturing and isolation of single pathogen colonies

Diagnostic samples (blood culture, pleural fluid, and respiratory material) obtained from the site of infection were directly processed at the medical microbiology unit at UHL. Respiratory samples and pleural fluid were incubated on agar media (blood agar, chocolate agar, MacConkey agar, Columbia CAP agar, and Sabouraud dextrose agar) and in liquid media under aerobic and anaerobic conditions at 37°C for 48 hours Blood cultures were processed using the BacTAlert System (Biomerieux, Lyon, France) and being incubated up to 7 days. Growth-positive blood cultures were streaked onto agar plates (Blood agar, chocolate agar, MacConkey agar, Columbia CAP agar, and Sabouraud dextrose agar) and incubated aerobically and anaerobically at 37°C. The pathogenic species were identified via MALDI-ToF (MALDI Biotyper, Bruker Daltonics) and antibiotic susceptibility testing was performed using broth dilution on one colony per pathogenic species. The evaluated isolate and up to nine additional colony isolates of each identified opportunistic pathogenic species’ morphology were stored in 30:70 glycerol/PBS in separate cryotubes at -80°C. After shipment of frozen isolates to the Max Planck Institute for Infection Biology, all isolates were cultured to validate their purity, re-identified by MALDI-ToF (VITEK MS, bioMérieux), and restreaked on ½ of a blood agar plate to produce sufficient biomass for whole-genome sequencing. One isolate of patient 21, expected to be *E. coli*, was identified as *Morganella morganii* and was excluded from further analysis.

For each confirmed HAI, we performed targeted culturing of the pathogen species within all microbiome specimens collected from that patient (**Figure S3**). Therefore, the microbiome sample and gastric aspirate sample of the HAI patients were thawed on ice, vortexed for 30 sec and 150 µl of serial ten-fold dilutions between 10^-2^ to 10^-7^ were plated on chromogenic or selective media plates (CHROMagar Acinetobacter [Mast Group; cat. no. 201481], Bacteroides bile esculine agar [NeoLab; cat. no. M805 and FD062], Mannitol salt agar [Thermo Fisher Scientific; cat. no. R453902], CHROMagar Orientation [Mast Group; cat. no. 201410], CHROMID *P. aeruginosa* agar [bioMérieux; cat. no. 43642], Simmons citrate agar [Thermo Fisher Scientific; cat. no. CM0155B] with inositol [Sigma-Aldrich; cat. no. I5125], Brilliance UTI Clarity agar [Thermo Fisher Scientific; cat. no. PO5159A], Stenotrophomonas selective agar [HiMedia; cat. no. M1965] with VIA supplement [HiMedia; cat. no. FD312]) using a semi-automated workflow set up on a TecanFluent liquid handling workstation equipped with the Scirobotics PetriLab hardware. To control for contamination during plating, a PBS control was taken along for each run, plated on Columbia blood agar (Carl ROTH; cat. no. X919.1) with 5% sheep blood (Thermo Fisher Scientific; cat. no. SR0051E) and incubated aerobically or anaerobically depending on the focal species. In addition, control air-swabs taken during microbiome sampling were plated undiluted. No growth was detected after up to 48 h for any control swab of the HAI patients. After incubation at 37°C between 24 h and 72 h, up to 20 single colonies of the pathogen species’ morphology were restreaked on a blood agar plate quadrant (*Proteus mirabilis* on half of a blood agar plate containing 3% agar) and incubated for another 24 h to 72 h at the same conditions to generate ∼ 100 mg biomass. Species of each isolate was confirmed using MALDI-ToF (VITEK MS, bioMérieux) and biomass was split after centrifugation at 16,500 g for 3 min in half for viable, long-term storage at -80°C (in 30:70 glycerin/PBS) and gDNA extraction.

### Library preparation and whole-genome sequencing

Cell pellets were lysed in a bead beater (Precellys Evolution, Bertin) using 400 mg glass beads (0.25-0.5 mm) in 180 µl B1 buffer ^61^ and 200 µl EB buffer at 7,200 rpm for 3×30 sec with 10 sec pause. Debris was removed at 12,500 g for 3 min. 25 µl Proteinase K (Macherey-Nagel; cat. no. 740506) was added and incubated at 56°C 450 rpm for 1 h. The lysate was used as input to the NucleoSpin Tissue Mini kit (Macherey-Nagel; cat. no. 740952.250S) following manufacturer’s instructions with self-made buffers exchanging the B3 buffer, BW buffer, B5 buffer, and BE buffer with the B2 buffer ^61^, PNI buffer ^61^, 80% ethanol, and 5 mM Tris-HCl (pH 8), respectively. DNA was quantified and normalized to 1 ng/µl using SYBR Green I (Thermo Fisher Scientific, cat. no. S7563) as described in ^62^.

Libraries were prepared on 96-well plates with alternating species on neighboring wells to allow identification of crosstalk between wells. Libraries were prepared using the Hackflex v1 protocol ^63^ with small modifications. For the tagmentation buffer, we used 20 mM Tris (pH 10.6), 10 mM MgCl_2_ and 20% Dimethylformamide adjusted to pH 7.6 with acetic acid. Amplification was done by using 11.1 µl KAPA HiFi HotStart Library Amp Kit (Roche; cat. no. KK2612), 2.2 µl of each i5 and i7 index primer (10 µM) respectively and 4.5 µl nuclease-free water. The PCR was performed using a two-step denaturation followed by 14 cycles (72°C for 3 min; 98°C for 5 min; 14 cycles with 98°C [10 sec] - 62°C [30 sec] - 72°C [30 sec]; 72°C for 5 min). Removal of large fragments were included for the library purifications and size selection. Therefore, 18 µl of the amplified library, 18 µl self-made 2% SPRI beads (2% SeraMag beads [Cytiva; cat. no. 65152105050250], 18% PEG-8000, 1 M NaCl, 10 mM Tris-HCl [pH 8.0], 1 mM EDTA [pH 8.0], 0.05% Tween-20) and 16.2 µl nuclease-free water were mixed and separated on a magnetic rack. 48 µl of the supernatant was mixed with 60 µl self-made 4% SPRI beads (as above with 4% SeraMag beads) and 52 µl nuclease-free water. Beads were washed twice with 200 µl freshly prepared 80% ethanol, dried for 5 min at room temperature and libraries were eluted in 32 µl resuspension buffer (10 mM Tris-HCl [pH 8.0], 0.05% Tween-20). Libraries were quantified using SYBR Green I as described in ^62^ and the quality of library preparation was assessed on a random subset for each species using a Fragment Analyzer (Agilent Technologies; DNF-915). Libraries were pooled equimolarly and sequenced at the Sequencing Core Facility of the Max Planck Institute for Molecular Genetics (Berlin, Germany) using either the NextSeq 500 (PE150) or NovaSeq 6000 (PE100) to reach ∼1.6 million paired reads per isolate.

### Classification of isolates into lineages

Single-isolate whole-genome sequences were clustered into lineages using core-genome SNV calls via an alignment-based approach using a custom snakemake ^64^ workflow following a previously published approach ^11^. Briefly, adapter sequences of raw reads were removed via cutadapt ^65^ (v 1.18) and low quality bases were trimmed using Sickle ^66^ (-q 20, -l 50 -x -n; v 1.33). Quality-filtered reads of each sequenced isolate was assessed for contamination introduced during high-throughput library preparation using Kraken2 ^67^ (v 2.0.8) and Bracken ^68^ (v 2.5) with a custom built database (including archaea, bacteria, fungi, human, plasmid, protozoa, UniVec, viral; build date: June 8th, 2022), a read length of 100 and a k-mer length of 35. This revealed an isolate-misclassification by MALDI-ToF of *Enterobacter cloacae* in patient 7 and *Klebsiella oxytoca* in patient 10 which have been identified to be *E. hormaechei* (both members of the *Enterobacter cloacae* complex ^69^) and *K. michiganensis* isolates (both members of the *Klebsiella oxytoca* complex ^70^), respectively. Further, four isolates indicative of cross-contamination (< 80% of reads supporting the focal species and > 10% of all reads classified as a species of another genus) and one isolate identified as different species (1.4% supported the focal species [*B. thetaiotaomicron*]; 68.82% of reads supporting another genus [*Parabacteroides*]) have been excluded from the subsequent analysis.

Based on the taxonomic classification, quality-filtered reads were aligned to their respective reference genomes (*Acinetobacter pittii:* GCF_000191145.1*, Bacteroides thetaiotaomicron:* GCF_014131755.1*, Escherichia coli:* GCF_000008865.2*, Enterobacter hormaechei:* GCF_000328885.1*, Klebsiella michiganensis:* GCF_015139575.1*, Pseudomonas aeruginosa:* GCF_000006765.1*, Proteus mirabilis:* GCF_000069965.1*, Staphylococcus aureus:* GCF_000013425.1*, Stenotrophomonas maltophilia:* GCF_900475405.1) using Bowtie2 ^71^ (-X 2000 --no-mixed --dovetail; v 2.2.6) and deduplicated using Picard MarkDuplicates ^72^ (v 2.26.11) with default parameters. Samtools ^73^ (v 1.5) mpileup (-q30 -x -s -O -d3000) was used to calculate genotype likelihoods at candidate variant sites (mapping quality ≥30) for every isolate. Candidate SNVs were subsequently called via bcftools ^73^ (v 1.2) call (-c) and filtered via bcftools view (-v snps, allele frequency [-q] ≥ 0.75).

We customized a previous pipeline protocol ^11^ which aggregates quality and summary metrics (strand-specific base calls, genotype confidence [FQ score], and coverage) for every variant position across all samples of the same species into multi-dimensional matrices enabling the quality assessment of each identified candidate SNV per pathogen-patient pair. First, isolates with a median genome-wide coverage below 8 indicative of insufficient genome resolution for high-confidence variant calling were removed from subsequent analysis (6/751). Second, we defined for every isolate the allele with the highest allele frequency at a given position as base call, if (1) the position had a major allele frequency of at least 0.95, (2) a genotype confidence (FQ score) below -30, (3) was supported by at least 2 reads per strand, and (4) less than 50% of reads at that position had an indel call in proximity of 3 nucleotides. Third, every candidate SNV was required to have a median coverage across all genomes of at least 4. Positions which failed to satisfy any filtering parameter, indicative of misalignments or insufficient evidence for high-confidence calls, were masked by setting the base call as ambiguous (N). Lastly, to remove co-segregating SNVs in close proximity which likely emerged from recombination, we calculated the Pearson correlation coefficient amongst filtered candidate SNVs per allele within a 1 kb window around every position and masked those which exceeded a correlation coefficient above 0.75. For lineage identification, we defined the core-genome SNVs as positions with less than 1% of ambiguous calls across all isolate genomes per pathogen-patient pair. A maximum-likelihood tree was computed using RAxML-ng ^74^ (v 1.2.2) with the GTR+G model. Branch lengths were rescaled using the Felsenstein ascertainment bias correction (ASC_FELS) integrated into the RAxML-ng suite with invariant sites defined as any monoallelic genomic position which would be considered for variant identification (median coverage ≥ 4, see thresholds above). Phylogenetic trees were rooted using as outgroup a reference genome polarizing the largest fraction of retained SNVs selected from a set of publicly available genomes of the species (**Table S4**) and visualized using ggtree ^75^ (v 3.14.0).

To further classify identified lineages, each isolate was assigned to a sequence type (ST) using SRST2 ^76^ (v 0.2.0) with default parameters where a species-specific MLST was available in pubMLST (https://pubmlst.org/) (**Table S5**). For *Escherichia coli* the Achtman scheme was used. STs of isolates with identified uncertainties (flagged with ‘?’) and mismatches (flagged with ‘*’) were considered valid when the suspected ST agreed with all other isolates of the lineage (6 cases), while not identifiable sequence types were turned to ‘NA’. To calculate a data-driven lineage cutoff within a species, we calculated the pairwise core-genome SNV distances for all isolates of each pathogen-patient pair. Separate lineages were identified to be further than 30 SNVs apart in agreement with the sequence typing and phylogenetic inference (**Figure 1c, Figure S13-S21**).

### Lineage-specific phylogenetic inference

To reconstruct the within-patient evolution of the pathogen species’ lineages genome-wide, including any isolate-specific accessory genome, we generated a lineage-wide *de novo* genome assembly for each pathogen-patient pair. To minimize contamination, we used only isolates with ≥ 80% of reads assigned to the focal species level during Bracken analysis (see “Classification of isolates into lineages”; **Table S5**). We randomly chose 250,000 reads per isolate that were phylogenetically assigned to a specific lineage (see “Classification of isolates into lineages”) to ensure equal representation of every isolate’s genetic composition. For pathogen lineages represented by only a single isolate (*E. coli* [ST131] from patient 7 and the minor lineage of *P. mirabilis* from patient 21) all sequencing reads available were used to maximise assembly quality. Assemblies have been generated using SPAdes ^77^ (v 3.15.4; --careful) and contigs longer than 500 bp whose assembly k-mer coverage was at least 10% of that of the largest contig within the assembly were retained. Completeness (99.99%-100%) and contamination (0.02%-0.86%) were evaluated via CheckM2 ^78^, indicative of high-quality genomes, and taxonomically confirmed via GTDB-Tk ^79^ (**Table S6**). Assemblies were annotated using Bakta ^80^ (v 1.6.1) with the full database (v 4.0) (**Table S6**). Moreover, for comparative convergent evolutionary analyses between pathogen species we clustered ortholog genes between assembled genomes using cd-hit ^81^ (v 4.8.1; -s 0.9 -c 0.95).

For variant identification, we aligned quality-filtered reads of all isolates of a species per patient (including those flagged due to Bracken species classification rates < 80%) against the *de novo* assembled lineage-specific genome using Bowtie2 (-X 2000 --no-mixed --dovetail; v 2.2.6) ^71^. To gain comprehensive insights about the body-wide diversity, we also included reads from four re-sequenced isolates which were confirmed to be contamination-free after previous exclusion due to indications of cross-contamination of the initial sequencing data (< 80% of reads supporting the focal species and > 10% of all reads classified as a species of another genus). Candidate SNVs were called as described above (see “Classification of isolates into lineages”), with one modification: to include the species accessory genome all sites were retained if the candidate position had a median coverage of at least 4 across samples with coverage at that position.

Phylogenetic trees per lineage were calculated using all retained SNVs and RAxML-ng ^74^ (v 1.2.2) with the GTR+G model. Branch lengths were rescaled using the Felsenstein ascertainment bias correction (ASC_FELS) integrated into the RAxML-ng suite with invariant sites defined as any monoallelic genomic position which is considered for potential variant identification (median coverage ≥ 4 across samples with coverage at that position). Phylogenetic trees were rooted using as an outgroup a reference genome polarizing the largest fraction of retained SNVs selected from a set of publicly available genomes of the species (**Table S4**). The phylogeny was visualized with ggtree ^75^ (v 3.14.0) if at least one SNV was observed (lineages without any SNV: *E. coli* ST131 [patient 7], *S. maltophilia* [patient 10], *P. aeruginosa* [patient 21], minor lineage of *P. mirabilis* [patient 21]) .

Muller plots were generated for *E. hormaechei* by considering all SNVs and indels for which we observed a substantial frequency change of at least 0.3 within the population over time. We aggregated genotype frequency shifts for all those SNVs and indels for all isolates combined as well as for isolates per microbiome niche individually. The frequencies were log-transformed to account for exponential growth and improve interpolation. Timepoints were removed if no isolates were collected. Piecewise Cubic Hermite Interpolating Polynomial (PCHIP) interpolation was performed to each genotype trajectory over time, respectively, using scipy ^82^ (v 1.15.2). The resulting back-transformed frequency trajectories were normalized to proportions per timepoint and visualized using ggmuller ^83^ (v 0.5.6).

### Lineage-specific indel analysis

Using the lineage-specific genome alignments, candidate indels were called jointly using freebayes ^84^ (v 1.3.6) in haploid mode and sites identified as insertion, deletion and complex were extracted. For every candidate indel summary metrics (indel sequence, indel support, length, genotype likelihood, and coverage) of the reference and alternative allele with the highest genotype likelihood have been collapsed into a multidimensional data structure for downstream filtering. To minimize false-positive calls, the candidate sites were filtered to exceed coverage ≥ 3 and a major allele frequency ≥ 80%. Sites were masked if the absolute genotype likelihood difference between reference and the best-scoring alternative allele was below 10. Moreover, positions without a major allele frequency ≥ 80% in at least 20% of samples - indicative for regions with misalignments - were removed. For *E. hormaechei*, identified indels were linked with quantified differences in antimicrobial resistance profiles. Moreover, multiple-sequence alignments of single isolate-specific assemblies were generated using SPAdes ^77^ (v 3.15.4; --careful) to confirm *ampD* truncations in *E. hormaechei*. To avoid false positive indel identifications due to assembly artefacts we aligned the reads of the respective isolate against the *de novo* assembled single isolate-specific genome via Bowtie2 ^71^ (-X 2000 --no-mixed --dovetail; v 2.2.6) and adjusted inconsistencies using pilon ^85^ (v 1.24). Contigs smaller than 500 bp as well as those whose assembly k-mer coverage was below 10% of the largest contig within the assembly suggestive of contamination due to poor read support were removed. Multiple sequence alignments (using MAFFT ^86^ [v 7.508]) for *ampD* sequences confirmed the observed *ampD* truncations and identified three additional *ampD* truncations (amino acid positions 93, 130, and 160) within 8 isolates (P07_1047, P07_1540, P07_1049, P07_1545, P07_1553, P07_1046, P07_1201, P07_1149) unobserved previously. Indels are annotated by their amino acid location to improve readability within graphics. For complete annotations see **Table S7**.

### Collector’s curve analysis

Collector’s curves for the DMRCA and SNVs have been calculated for every observed lineage per patient, timepoint and location (site of infection, collapsed microbiome sites [nasal, oral, rectal, skin], gastric samples) with at least five isolates. For each timepoint the number of collected isolates (*n*) was randomly sampled with replacement to subset sizes *x* (1 ≤ *x* ≤ *n*) with 100 replicates per subset size. For each replicate subset, the mean distance to their subset’s most recent common ancestor (DMRCA) and the number of unique SNVs were calculated. The ancestral state of each position was inferred based on a rooted parsimonious tree generated by PHYLIP’s dnapars ^87^ (v 3.695) using TreeTime’s ^88^ (v 0.9.2) ancestral module. The mean and standard deviation across replicates of each subset size *x* were calculated and reported.

### Inference of colonization timings

To obtain species-specific molecular clocks, the ancestral sequence was inferred for each pathogen lineage using a rooted parsimonious tree generated by PHYLIP’s dnapars ^87^ (v 3.695) and TreeTime’s ^88^ (v 0.9.2) ancestral module. The inferred ancestral sequence was used to calculate the distance to the most recent common ancestor (DMRCA) for every microbiome isolate, respectively, and normalized using the number of genomic sites exceeding the minimum coverage required to identify a variable site per isolate (≥ 4 reads). The molecular clock was inferred using linear regression of all normalized DMRCA counts over time per lineage using scipy ^82^ (v 1.15.2). We only observed for *E. hormaechei* a positive correlation (r^2^ = 0.347; **Figure S15**).

We validated the applicability of published molecular clocks by comparing the mutational spectrum observed within the patients’ pathogen lineage to the mutational spectrum between publicly available genomes of the species (**Table S4**). All SNVs (filter criteria as described in “Classification of isolates into lineages”) were categorized into transitions and transversions. The observed transition and transversion ratio among the publicly available reference genomes and within the pathogen lineage were compared using a χ^2^ test. One pathogen lineage (*E. coli* ST 23 from patient 21) showed a significant deviation from the expected spectrum and was excluded from further analysis (**Figure S16**).

For the remaining pathogen lineages, we only observed published molecular clocks for *S. aureus* ^11,26,89–97^, among which we identified a study reporting a sequence-type-specific molecular clock which was reanalysed using their published criteria ^11^. For both remaining pathogen species with a lineage- (*E. hormaechei*) or sequence-type-specific (*S. aureus*) molecular clock, the time to the most recent common ancestor (TMRCA) was estimated using the mean DMRCA of the first microbiome sampling timepoint with at least 8 isolates (always before or within 6 h of HAI onset). To propagate the error of the interval estimates of the molecular clocks and the observed overall low mutation counts we fitted a gamma distribution to the molecular clock parameters (α =(molecular clock)^2^/(variance of molecular clock); θ = (variance of molecular clock)/(molecular clock)) by minimizing the sum of squares between the estimated and given log-transformed parameters using the L-BFGS-B algorithm within the scipy’s optimize module ^82^ (v 1.15.2). We jointly sampled 10,000 values from these gamma distributions and bootstrapped DMRCA counts to estimate the confidence intervals of the TMRCAs.

### Within-lineage mobile genetic elements

Mobile genetic elements (MGE) were inferred within each observed lineage following a previously published approach ^11^. Briefly, using the lineage-specific *de novo* assemblies we identify regions missing in at least one isolate using a z-score normalizing coverage at each position first within an isolate and, second, across isolates. MGE-candidate regions, called per sample, needed to be longer than 750 bp and the double z-score normalized value for every position within that region needed to be below -0.5 with a mean value below -1 across the entire region, indicative of consistent coverage suppression at that region. Candidate regions had to be supported by one isolate with a median coverage of 0 (absence) as well as one isolate with a median coverage above 5 (presence). All identified MGEs across samples were merged into a consecutive MGE if they overlapped with at least 10 bp (**Table S7**).

### Analysis of public *E. hormaechei* genomes

Publicly available whole-genome sequences of *E. hormaechei* isolates were used to assess the frequency of *fimZ* mutations in the clinical context on a global-scale. Therefore, we downloaded *Enterobacter* paired-end Illumina whole-genome sequences from the Pathogen Detection database (accession date 17.12.2025; search term on the Isolate Browser: “Enterobacter hormaechei”) ^98^ as well as the CDC HAI-Seq Gram-negative bacteria BioProject (PRJNA288601, accession date 09.11.2023) ^99^ from respiratory samples of the human host. Whole-genome sequences were evaluated to be *E. hormaechei* (see “Classification of isolates into lineages”) and 491 genomes retained stemming from a total of 42 BioProjects (**Table S3**). Reads were aligned against the *de novo* assembled *E. hormaechei* genome of patient 7, samples with median coverage < 8 - indicative of insufficient genome resolution for high-confidence variant calling - were removed (11/491 genomes), and core-genome SNVs were identified and filtered as described before except the removal of co-segregating SNVs (see “Classification of isolates into lineages”). Instead, genomic regions with evidence for recombination events were identified using Gubbins ^100^ (v 3.4) with default parameters and masked. This retained a total of 79,610 SNVs. To assess signatures of molecular evolution we calculated the dN/dS-scores for every mutated gene (n = 2,909), considering the observed genome-wide codon usage and mutational spectrum. The phylogeny was reconstructed with RAxML-ng ^74^ (v 1.2.2) using the GTR+G model, with the topological stability of the phylogeny assessed using the Transfer Bootstrap Expectation ^101^ values derived from 1,000 bootstraps. The phylogeny was visualized using ggtree ^75^ (v 3.14.0).

### Protein structure prediction

To estimate the location of the mutation, we predicted the 3D protein structure of FimZ using the wildtype variant as input for AlphaFold 3 ^102^ (v 3.0.0) using default options to generate 5 models. The highest-scoring model (ranking score: 0.82; mean pLDDT: 85.7) has been used for subsequent analysis. The structure was converted to a PQR file using gemmi convert ^103^ (v 0.7.1) and PDB2PQR ^104^ (v 3.7.1) using the PARSE force field at pH 7. The electrostatic potential was calculated using APBS ^104^ (v 3.4.1) in mg-auto mode and default options. The grid has been defined using the psize.py script provided with APBS. The structure and electrostatic potential was visualized in UCSF ChimeraX ^105^ (v 1.6.1) centering the color scale on the mean electrostatic potential with the maximum colors corresponding to ±3.

### Transcriptional expression analysis (Real-time qPCR)

*E. hormaechei* cultures of different growth stages (early- [OD_600_ 0.1] and late-log [OD_600_ 1]) were harvested at 7,500 g for 10 min. 5*10^8^ CFU were resuspended in 750 µl TRIzol (Thermo Fisher Scientific; cat. no. 15596018), vortexed vigorously, and incubated for 5 min at room temperature. Samples were mixed with 200 µl chloroform, inverted 20 times, and incubated for 3 min at room temperature. Phases were separated at 12,000 g for 15 min at 4°C. Aqueous phase was mixed 1:1 with 99% ethanol and total RNA extraction was proceeded using the Direct-zol RNA Miniprep Kit (Zymo Research, cat. no. R2050) following the manufacturer’s instructions including the DNase I digestion. Total RNA was quantified using the NanoDrop ND-1000 spectrophotometer and cDNA synthesis was done with 250 ng total RNA using the High Capacity cDNA Reverse Transcription Kit (Applied Biosystems, cat. no. 4368814) with random hexamer primers. RT-qPCR was performed targeting the genes *rpoD*, *fimZ*, and *fimA* (**Table S8**) using the iTaq Universal SYBR Green Supermix (Bio-Rad, cat. no. 1725121) in a qTOWER^3^. gDNA levels were estimated using reverse transcriptase-negative controls and samples with gDNA levels ≥ 3.125% (ΔCt ≥ 5) were removed to ensure accurate estimation of the gene expressions ^106^. Remaining samples were corrected for their gDNA levels and normalized to the housekeeping gene *rpoD*. The relative expression was calculated using the ΔΔCt method ^107^.

### Growth curve assays

Overnight cultures of *E. hormaechei* isolates were diluted in LB media to OD_600_ 0.01 and seeded into a transparent flat-bottom 96-well plate with 3 technical replicates per isolate. Growth at 37°C and 142 rpm was measured semi-automatically every 20 min via the OD_595_ using the Tecan Infinite 200 Pro M Nano+. Optical density values were adjusted using the average optical density measurement of abiotic medium at every respective timepoint and growth curves were smoothed in a rolling window of 40 min.

### Scanning electron microscopy

*E. hormaechei* isolates were grown to late-log (OD_600_ 1; growth stages mid- [OD_600_ 0.4] and stationary [overnight culture] showed similar results but are not shown). 2.5*10^8^ CFU were harvested at 500 g for 10 min, washed with 1 ml PBS and fixed in 2.5% glutaraldehyde. Subsequently, fixed bacteria were added directly onto poly-L-lysine-treated glass coverslips (Carl ROTH; cat. no. P231.1). The samples were post-fixed in 0.5% osmium-tetroxide, tannic acid and osmium-tetroxide for 15 min, respectively. A graded ethanol series was applied to dehydrate the coverslips and dried in carbon dioxide at critical point before they were vacuum coated with 3 nm carbon-platinum. Imaging was performed using a Zeiss LEO 1550 scanning electron microscope using an in-lens detector at 20 kV acceleration voltage.

### Antimicrobial susceptibility testing

Antimicrobial susceptibility testing was performed according to EUCAST guidelines and minimum inhibitory concentrations (MIC) were interpreted following the EUCAST Clinical Breakpoints Tables (v15.0) ^108^. In brief, selected *E. hormaechei* isolates were grown in Mueller-Hinton media. Cultures were diluted to 1.5*10^8^ CFU/ml and 100 µl were plated on Mueller-Hinton agar. Subsequently, ETEST (bioMérieux) of antibiotics, with which the respective patient had been treated, were applied, except for erythromycin, vancomycin, and ceftazidime (patient 7) due to following reasons: erythromycin was administered to stimulate gut activity rather than as an antibiotic and initial screening revealed high resistance (≥ 128 µg/ml), vancomycin specifically targets gram-positive bacteria but *E. hormaechei* is a gram-negative bacteria, and ceftazidime is a 3^rd^ cephalosporine like cefotaxime which revealed similar resistance profiles after an initial screen..

### Crystal violet assay

*E. hormaechei* isolates were inoculated in 96-well plates in λ broth at 30°C for 6 hours before 125 µl of 0.1% (wt/vol) crystal violet solution were dispensed into each well, followed by an incubation period of 45 min at room temperature. Next, the wells were washed twice with water to eliminate excess crystal violet, after which the plate was dried at 37°C overnight. 150 µl destaining solution (45% methanol, 45% water, and 10% glacial acetic acid) was added to each well, and incubated for 5 minutes at 25°C. Following incubation, 100 µl from each well was transferred to a 96-well flat bottom polystyrene plate and the OD_550_ was measured using a spectrophotometer. Isolates were grouped based on their *fimZ* genotype and analysed for statistical significance using a Mann-Whitney U test.

### Microfluidic biovolume measurement

Overnight cultures of *E. hormaechei* isolates back-diluted to an OD_600_ of 1.0 were inoculated into microfluidic chambers. After a 45-min incubation to allow for surface attachment, M9 minimal media with 0.5% glucose and Syto 9 stain was continuously flown into the chamber at a rate of 0.1 μl/min. Experiments were carried out at room temperature (22°C). Biofilms were incubated for 24 h prior to imaging. Images were quantified for biovolume with BiofilmQ ^109^. Isolates were grouped based on their *fimZ* genotype and analysed for statistical significance using a Mann-Whitney U test.

### Lung epithelial adherence assay

To measure the adherence to lung epithelial cells, 5.5*10^5^ Calu-3 cells (ATCC HTB-55) were seeded in a 24-well plate in 1 ml advanced DMEM (Thermo Fisher Scientific; cat. no. 12491015) containing 10% FBS (Thermo Fisher Scientific; cat. no. A5256801) and 1x Glutamax (Thermo Fisher Scientific; cat. no. 35050061). Cells were incubated at 37°C with 5% CO_2_ overnight before confluency was confirmed to be at 95% using an inverted microscope. Overnight cultures of *E. hormaechei* isolates were harvested by 7,500 g for 10 min, washed in 1 ml PBS and centrifuged again. The pellet was resuspended in advanced DMEM containing 10% FBS and 1x Glutamax to reach a density of 3*10^8^ CFU/ml and bacterial load was confirmed via CFU measurements. Calu-3 cells were carefully washed with 1 ml PBS before adding 1 ml of bacterial culture (multiplicity of infection: 1:20). To evaluate background adherence, an empty well has been seeded with 1 ml bacterial culture as well. The 24-well plate was centrifuged at 300 g for 5 min and incubated for 2 h at 37°C with 5% CO_2_. After incubation, wells were carefully washed 5 times with 1 ml PBS. Calu-3 cells with adherent bacteria were detached with 200 µl 0.025% Trypsin-EDTA (Thermo Fisher Scientific; cat. no. 25200056) for 20 min at 37°C and 5% CO_2_. 800 µl PBS was added and cells were transferred to a fresh 1.5 ml tube. After centrifugation at 7,500 g for 10 min, the pellet was resuspended thoroughly in LB media. *E. hormaechei* CFUs were determined via plating of several ten-fold dilutions. Background adherence of bacteria to the wells was removed using the average adherence of bacteria to wells not seeded with Calu-3 cells. To calculate the relative adherence, CFUs have been normalized to the measured bacterial load used as input for the adherence assay.

### *Drosophila* infection model

To evaluate the virulence of the identified *fimZ* [F126L] genotype, we used a systemic infection model of *Drosophila melanogaster* flies. *Drosophila* stocks (DrosDel *w*^1118^ iso and *Relish^E^*^20^ iso) were maintained at 25°C as described in ^110^. Frozen stocks of *E. hormaechei* isolates (wildtype: P07_2048, P07_2055; *fimZ* [F126L]: P07_2052, L00071_01_1) were streaked onto blood agar plates, inoculated at 37°C overnight, and one colony per isolate was inoculated in 50 ml LB at 37°C at 175 rpm overnight. Cultures were harvested at 1,500 g for 15 min at room temperature. Cell pellets were resuspended thoroughly in PBS to avoid any cell aggregation and subsequently diluted to OD_600_ 5. Systemic infections were performed by anaesthetizing 3-7 d old male DrosDel *w*^1118^ iso (wildtype) and *Relish^E^*^20^ iso flies with CO_2_ and pricking the thorax with a 0.15 mm minutien pin (Fine Science Tools; cat. no. 26002-15) mounted on a metal holder which has been immersed into the bacterial suspension. 20 infected flies were combined per vial and housed at 29°C with a 12:12 h light-dark cycle. The flies which did not recover within the first 2 h post infection were removed from the assay. Remaining flies were counted regularly up to 15 h or 168 h for *Relish^E^*^20^ iso and wildtype flies, respectively, with fresh food provided every 3-4 d.

To ensure comparable bacterial inoculum and assess the bacterial load of *Relish^E^*^20^ iso flies before death onset, CFUs of five individual flies per bacterial isolate and fly genotype were measured at 0 h and 10 h post infection, the start of increased hazard rate for *Relish^E^*^20^ iso flies. Therefore, flies were anesthetized, washed with 70% ethanol and subsequently homogenized with the Precellys Evolution at 7,200 rpm for 30 s in 500 µl PBS. Several ten-fold dilutions in PBS were generated and plated in triplicates on BHI plates. Following an incubation at 37°C overnight, the CFU were determined. Starting inocula were compared between bacterial isolates per experimental date and fly genotype to avoid biased results due to variance during fly infection. The CFU and survival assay was repeated three independent times with 50 recovered flies per isolate and fly genotype. We identified significant differences (Mann-Whitney U test, p ≤ 0.05) in the bacterial inoculum between bacterial wildtype and *fimZ* mutant genotype. Variance in bacterial inoculum led to the removal of one isolate (L00071_01_1 [*fimZ* [F126L]]) from the first experiment (wildtype and *Relish^E^*^20^ iso flies), the exclusion of the second experiment (*Relish^E^*^20^ iso flies) and one isolate (P07_2048 [wildtype]) from the second experiment (wildtype flies) from further analysis (**Figure S22a**, **Figure S23a**).

The survival efficiency of flies infected with *fimZ* [F126L] vs wildtype isolates was analysed in python using the lifelines module ^111^ (v 0.27.8). Survival curves per genotype were fitted using the Cox-proportional hazard model, with experimental dates included as covariates, and statistical difference assessed using the integrated Wald test.

### Validation of genotypes used for phenotypic assays

The genotypes of all stock isolates used for phenotypic assays have been confirmed by re-sequencing. Therefore, gDNA was extracted from isolates re-streaked from frozen long-term storage stocks and libraries were prepared as described above (see “DNA extraction and single-colony sequencing”) and sequenced to gain ∼ 0.5 Mio reads per isolate (Illuminia NovaSeq 6000 [PE100]). Obtained reads were aligned against their respective lineage-specific assembled genome after quality-filtering and SNVs as well as indels were called as described previously (see “Lineage-specific phylogenetic inference” and “Lineage-specific indel analysis”) with the adjustments of requiring a median genome coverage ≥2, one read on each strand and a minimal total coverage of 3 to account for the overall lower sequencing depth. This analysis revealed no new SNVs nor indels, validating the observed genotypes used for phenotyping.

## Data availability

Isolate genome data has been deposited in the European Nucleotide Archive under the accession PRJEB98220. Individual isolates alongside metadata and accession identifier are presented in **Table S5**. The complete code base alongside the annotated assembled lineage-specific genomes is available at https://github.com/fm-key-lab/HAI_evo_publication.

## Acknowledgements

We like to thank the past and present members of the Key lab for helpful discussion, Christian Goosmann & Volker Brinkmann from the MPIIB for their support with scanning electron microscopy, and Diane Schad (MPIIB) for support during graphic illustrations. At the University hospital Leipzig we would like to thank Annett Hennig-Rolle, Lisa Müller, the nurses and physicians for sample processing as well as the patients for participation. We thank Andreas Diefenbach for access to the VITEK MS at Labor Berlin. This work was supported by the Max Planck Society (to M.F., A.H., M.S., L.O., E.C., F.S., I.I., F.M.K.).

## Author contributions

Conceptualization and design of clinical cohort: B.P. and F.M.K.

Enrolled patients and collected clinical samples: B.P. and S.P.

Identified and isolated bacteria from clinical specimens: C-S.S. and N.L.

Isolated bacteria from microbiome specimens, extracted DNA, and prepared genomic libraries: M.F. and M.S.

Performed genomic analysis and interpretation: M.F. and F.M.K.

Performed in vitro and in vivo assays and interpretation: M.F., A.H., J.B.W., M.S., L.O., E.C., F.S., C.D., I.I., F.M.K.

Writing - Original Draft: M.F and F.M.K.

Writing - Review & Editing: M.F, A.H., J.B.W., S.P., N.P., C.D.N., I.I., B.P. and F.M.K.

