## Supplementary Figures for "Colonization, translocation, and evolution of opportunistic pathogens during healthcare-associated infections"

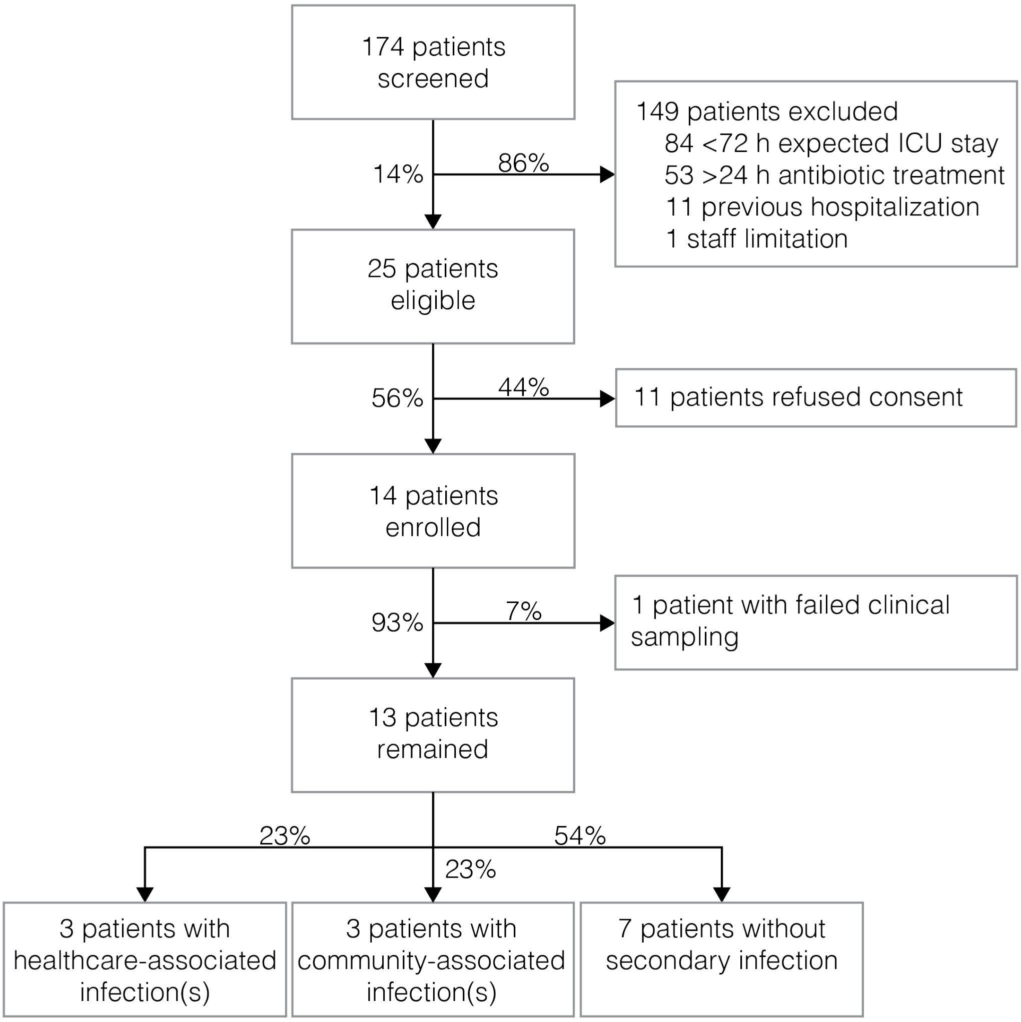


**Figure S1: Prospective screening of critically ill patients at the medical intensive care unit of the University Hospital Leipzig.**

Prospective screening of 174 critically ill patients within 3.5 months revealed 25 eligible patients which were expected to have an ICU stay exceeding 72 h, did not receive antibiotics within the last 30 days nor have been under treatment for more than 24 h, and were not hospitalized within the last 30 days. Of those, 14 patients consented to participate in this study, but one patient was excluded due to failure to retain the only clinical sample at healthcare-associated infection (HAI) onset. From 13 patients we observed HAIs in three patients. Percentages represent the proportion of patients classified in the proceeding step.

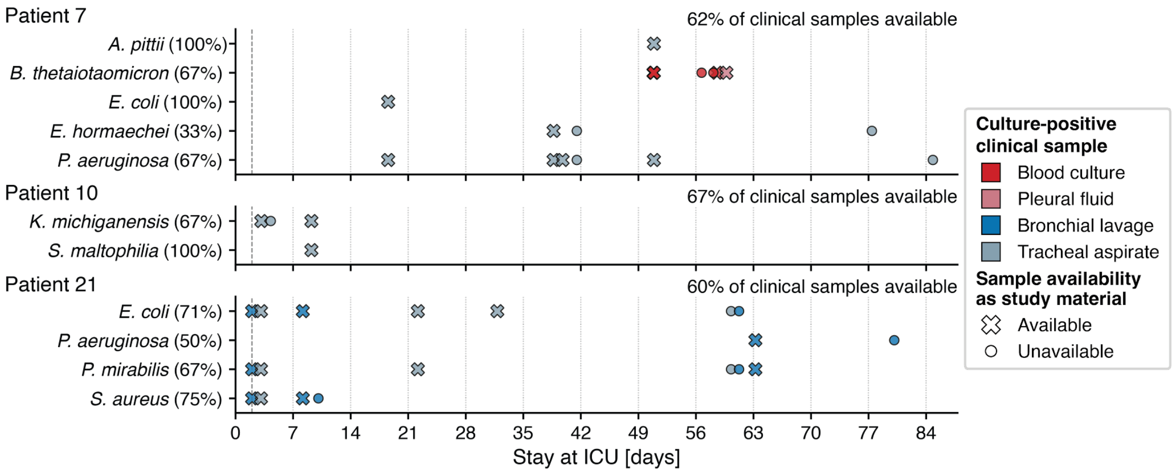


**Figure S2: Timepoints of healthcare-associated bloodstream and pneumonia diagnosis and successful collection of isolates from diagnostic patient material.**
Upon clinical suspicion of a HAI (see Methods), corresponding patient material was cultured at the medical microbiology unit of the University Hospital Leipzig. In case of *B. thetaiotaomicron* in patient 7, a contextual sample from the pleural empyema was taken during surgery. All culture-positive samples are shown, which confirmed a suspected HAI (> 48 h post admission, represented by a dashed line). However, due to logistical challenges during COVID19 and staff shortages, collection of isolates was only possible for a subset of the culture-positive specimens (crosses). The collection frequency per pathogen and patient is stated behind each species name and the overall frequency per patient is stated on top of each panel.

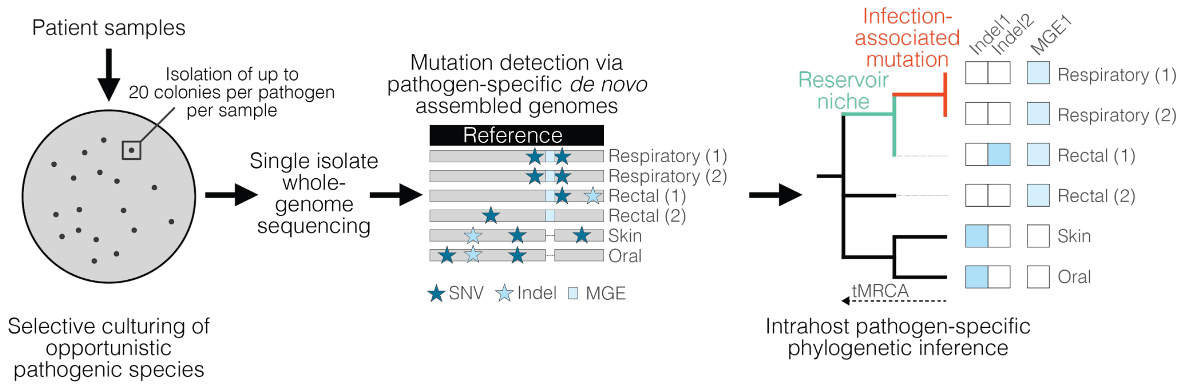


**Figure S3: Specimen processing from HAI patients to reconstruct the spatiotemporal pathogen-specific diversity.**

For every observed HAI, we performed targeted selective culturing of the identified pathogenic species within the patient's microbiome and collected up to 20 isolates per microbiome niche and time point. Whole-genome sequencing of all collected microbiome and clinical isolates was performed. Within-patient lineage-specific genomes were assembled and genetic variants (single nucleotide variants [SNV], indels and mobile genetic elements [MGE]) were called to identify possibly pathoadaptive signatures.

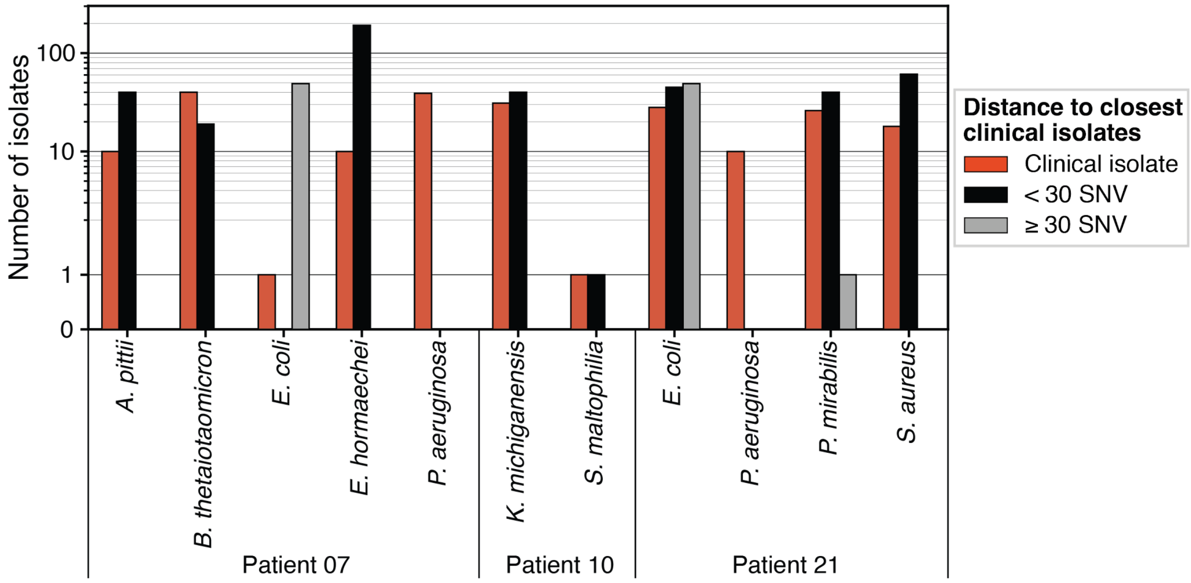


**Figure S4: Isolate collection per pathogen-patient pair.**
Isolate genomes are grouped: (1) isolates from the site of infection (clinical isolate), (2) microbiome isolates from the same lineage as a clinical isolate (< 30 SNV), and (3) microbiome isolates from a different lineage as the clinical isolates (≥ 30 SNV) (see **Figure 1c** and Methods “Classification of isolates into lineages”). Isolates from gastric samples are shown as clinical isolates (*K. michiganensis* [n = 20]). Pathogens are grouped on the x-axis per patient. The y-axis is in log-scale to improve visualization.

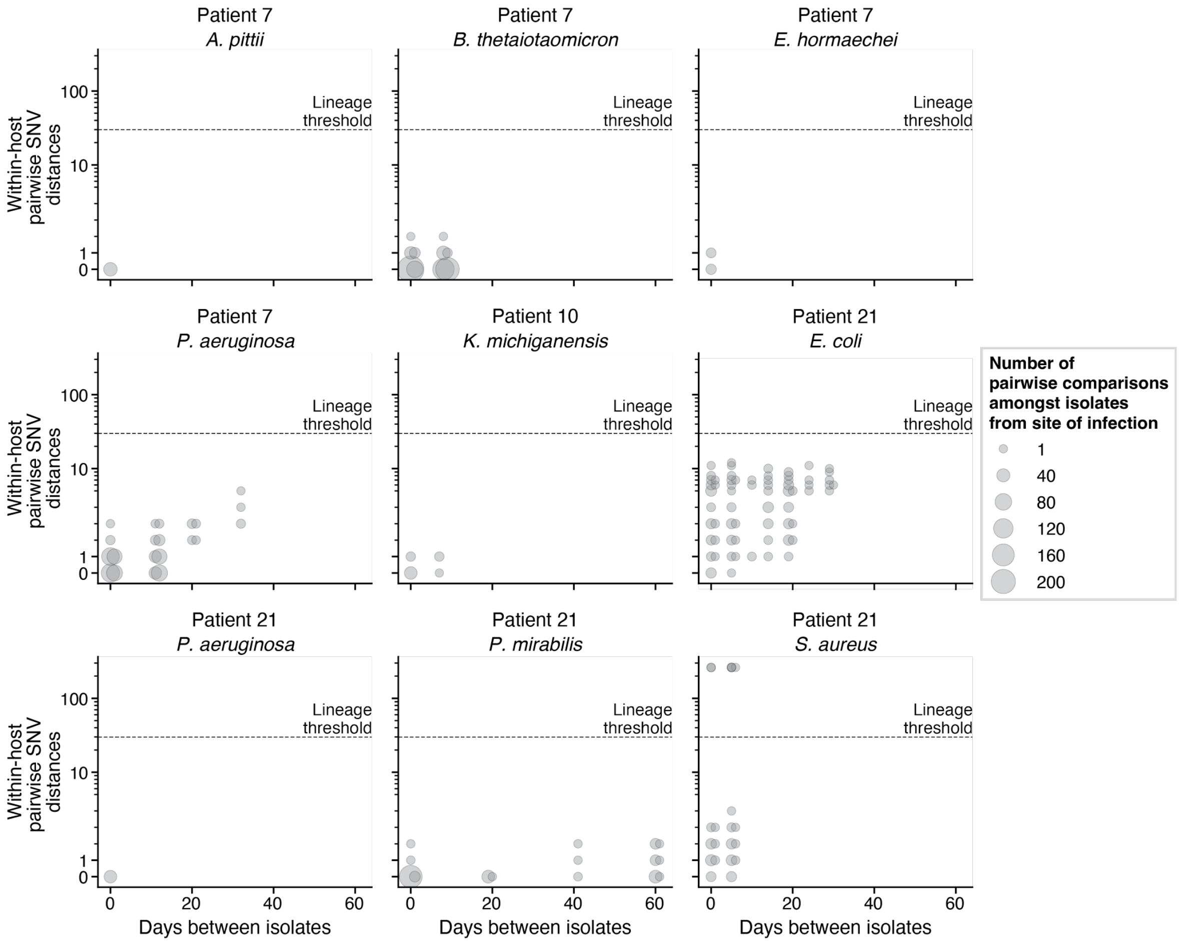


**Figure S5: Within-host pairwise SNV distances of clinical isolates reveal individual lineages propagate at the site of infection over time.**
Per pathogen-patient pair with at least two clinical isolates, the pairwise core-genome SNV distances were calculated across all collected clinical isolates from the site of infection over time. The number of pairwise comparisons are indicated through the size of the marker. The dashed line represents the data-derived threshold to cluster isolates into separate lineages (30 SNVs; **Figure 1c**). The y-axis is in log-scale to improve visualization.

9

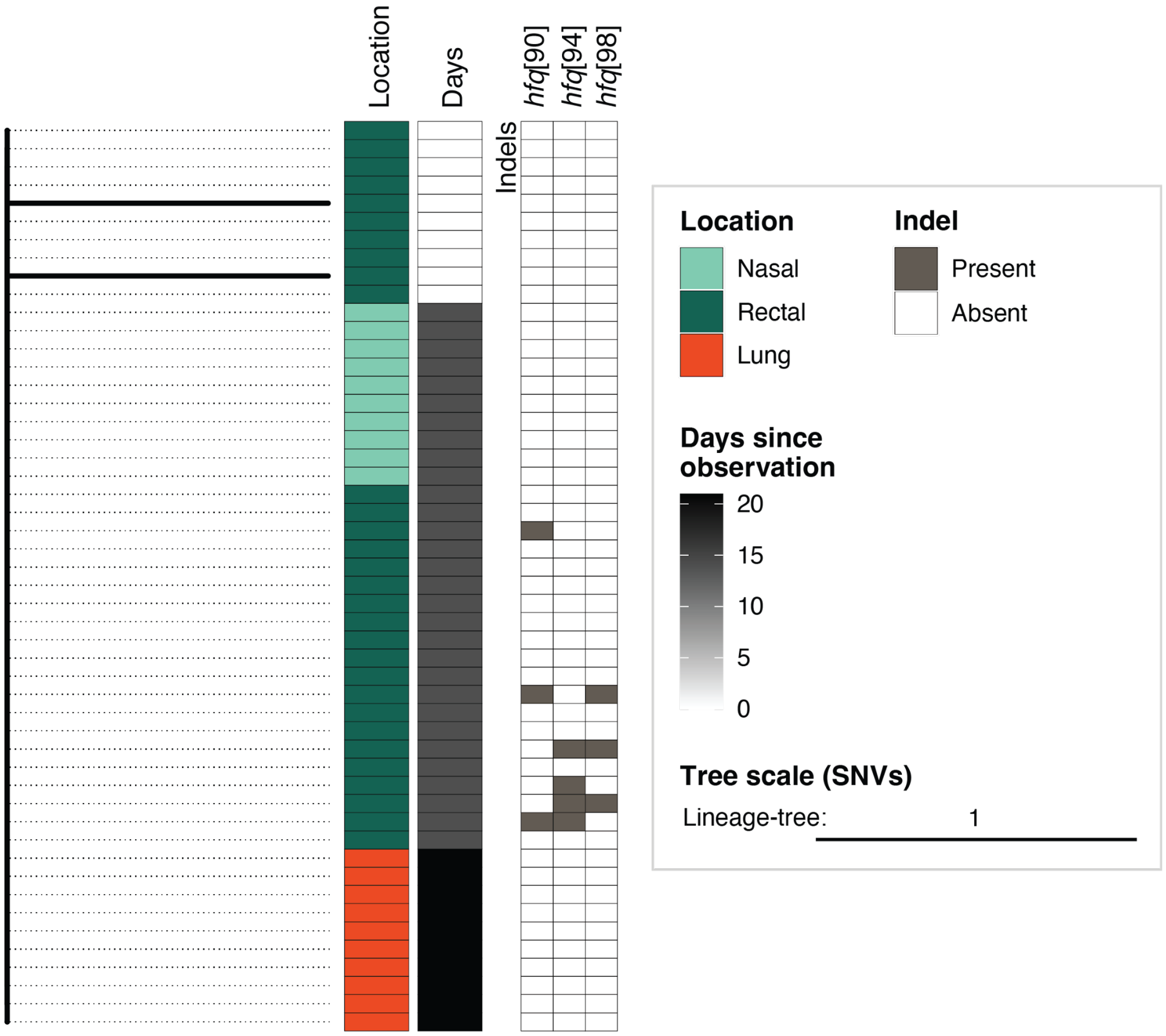


**Figure S6: Population structure of *Acinetobacter pittii* within patient 7.**
Maximum-likelihood phylogeny of the 50 *A. pittii* isolates from patient 7 using the lineage-specific alignment. The tree, based on two SNVs, is rooted on GCF_001577285.1 and the tree scale bar represents one SNV. The microbiome isolation source is shown in shades of green or red for isolates collected from the site of infection. Timepoints since first isolation (in days) are shown in grey scale. Non-singleton indels are shown in grey within a separate heatmap aligned to the tips of the tree with the gene and position of the mutated amino acid in squared brackets. To reduce complexity, mobile genetic elements are filtered to be supported by at least two isolates (≥ 95% coverage breadth) and lacking in two other isolates (≤ 75% coverage breadth) of which none was identified. All identified mutations of *A. pittii* can be found in **Table S7**.

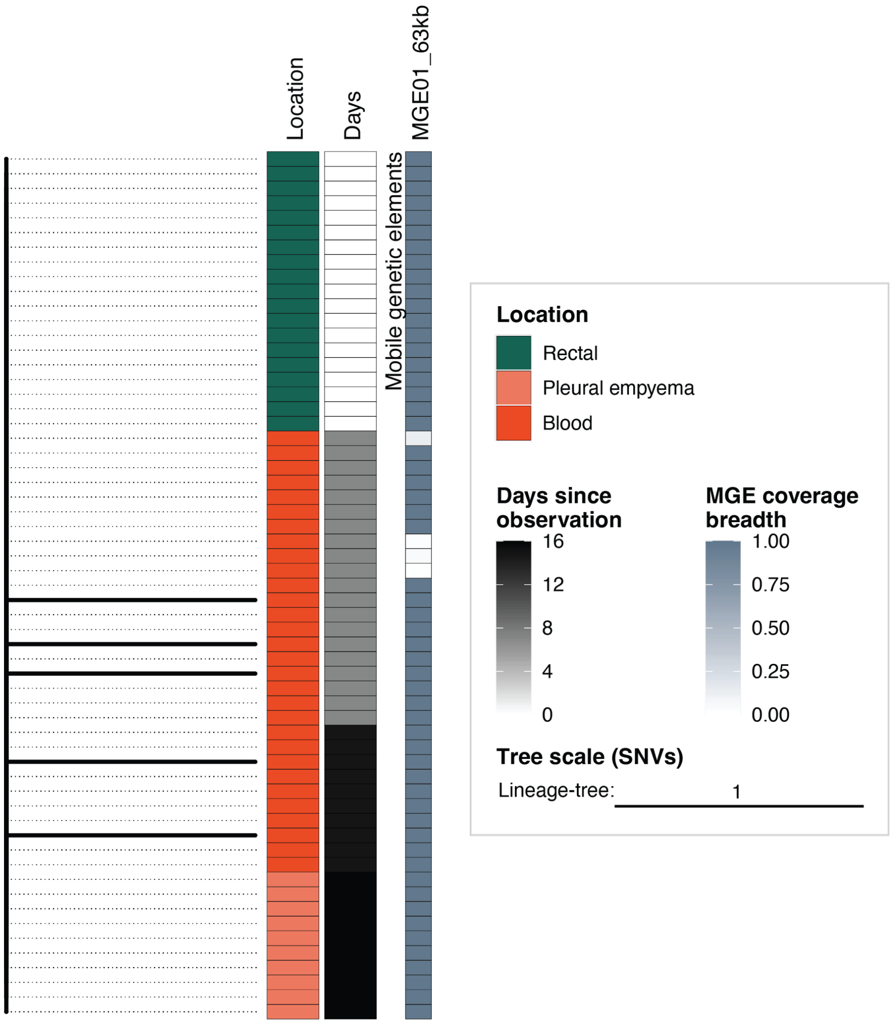


**Figure S7: Population structure of *Bacteroides thetaiotaomicron* within patient 7.** Maximum-likelihood phylogeny of the 59 *B. thetaiotaomicron* isolates from patient 7 using the lineage-specific alignment. The tree, based on five SNVs, is rooted on GCF_001314975.1 and the tree scale bar represents one SNV. The microbiome isolation source is shown in shades of green or red for isolates collected from the site of infection. Timepoints since first isolation (in days) are shown in grey scale. To reduce complexity, only mobile genetic elements supported by at least two isolates (≥ 95% coverage breadth) and lacking in two other isolates (≤ 75% coverage breadth) are shown by coverage breadth within a separate heatmap aligned to the tips of the tree. The MGEs are numbered consecutively and the length of the MGE is stated. No non-singleton indels have been observed. All identified mutations of *B. thetaiotaomicron* can be found in **Table S7**.

12

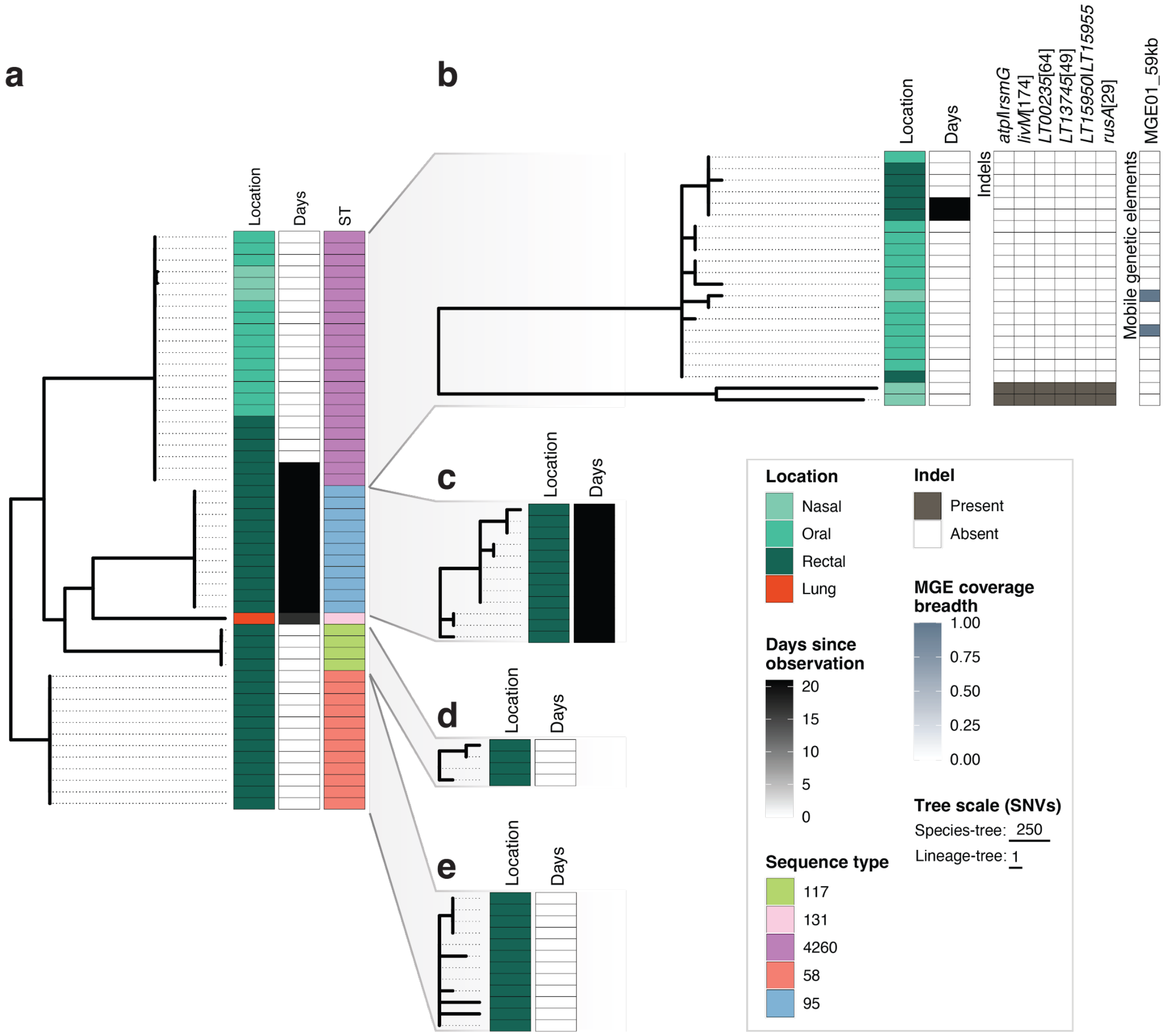


**Figure S8: Population structure of *Escherichia coli* within patient 7.**
**a.** Maximum-likelihood phylogeny of the 50 *E. coli* isolates from patient 7 using the core-genome alignment to show the entire population. The tree is rooted on GCF_000782595.1 and the respective species-specific tree scale bar represents 250 SNVs. The microbiome isolation source is shown in shades of green or red for isolates collected from the site of infection. Timepoints since first isolation (in days) are shown in grey scale. The sequence types (ST) per isolates are aligned to the tips of the tree.
**b-e.** Maximum-likelihood phylogeny per lineage excluding sequence type 131 with only one observed isolate. Phylogenies have been reconstructed using the lineage-specific alignment and rooted on GCF_008632595.1 (**b**), GCF_004380435.1 (**c**), GCF_000408225.1 (**d**), and GCF_001900555.1 (**e**). The trees are based on 72 (**b**), nine (**c**), four (**d**), and 10 (**e**) SNVs, respectively, scaled to the lineage-specific scale bar representing one SNV. Isolation timepoints and source use the same color scheme as in (**a**). For ST4260 (**b**), non-singleton indels are shown in grey within a separate heatmap aligned to the tips of the tree with the gene and position of the mutated amino acid in squared brackets, while for intergenic indels, adjacent genes are stated, separated by "|". To reduce complexity, only mobile genetic elements supported by at least two isolates (≥ 95% coverage breadth) and lacking in two other isolates (≤ 75% coverage breadth) are shown (one in ST4260) by coverage breadth within a separate heatmap aligned to the tips of the tree. The MGEs are numbered consecutively and the length of the MGE is stated. No mobile genetic elements (≥ 2 isolates ≥ 95% coverage breadth; ≥ 2 isolates ≤ 75% coverage breadth) nor non-singleton indels have been observed for the remaining lineages. All identified mutations within each lineage for *E. coli* from patient 7 can be found in **Table S7**.

14

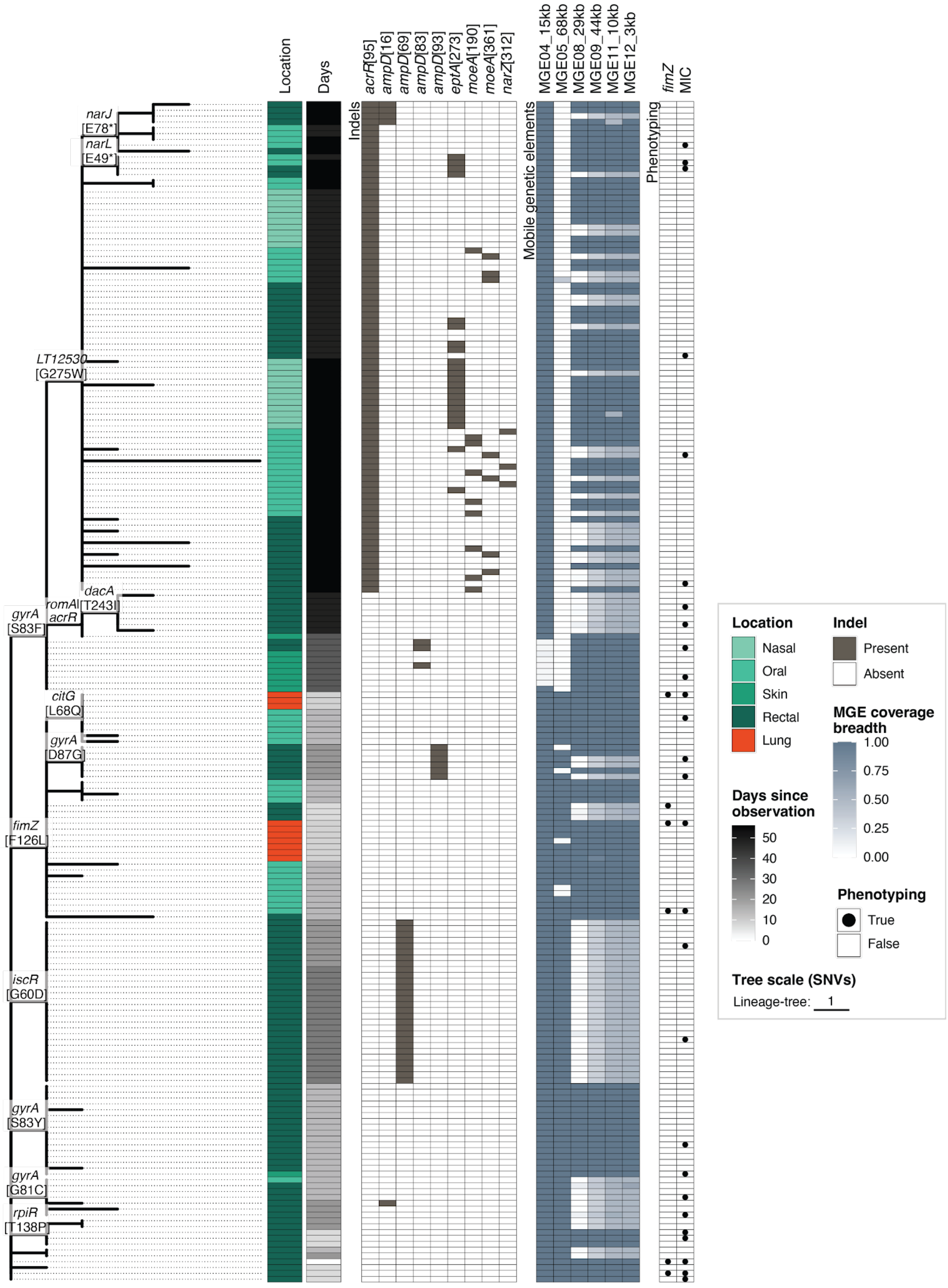


**Figure S9: Population structure of *Enterobacter hormaechei* within patient 7.**
Maximum-likelihood phylogeny as shown in **Figure 3a** with additional metadata. In brief, the phylogenetic relationship among the 202 *E. hormaechei* isolates from patient 7 is shown based on the 61 SNVs identified using the lineage-specific alignment. The tree is rooted on GCF_003408555.1 and the tree scale bar represents one SNV. The microbiome isolation source is shown in shades of

green or red for isolates collected from the site of infection. Timepoints since first isolation (in days) are shown in grey scale. Non-singleton indels are shown in grey within a separate heatmap aligned to the tips of the tree with the gene and position of the mutated amino acid in squared brackets. Mobile genetic elements are shown by coverage breadth within a separate heatmap aligned to the tips of the tree if it is supported by at least two isolates (≥ 95% coverage breadth) and lacking in two other isolates (≤ 75% coverage breadth) to reduce complexity. The MGEs are numbered consecutively and the length of the MGE is stated. Isolates used for phenotypic analysis are highlighted by black dots in the last heatmap. SNVs carried by those isolates are annotated on the branches within the tree. All identified mutations of *E. hormaechei* can be found in **Table S7**.

16

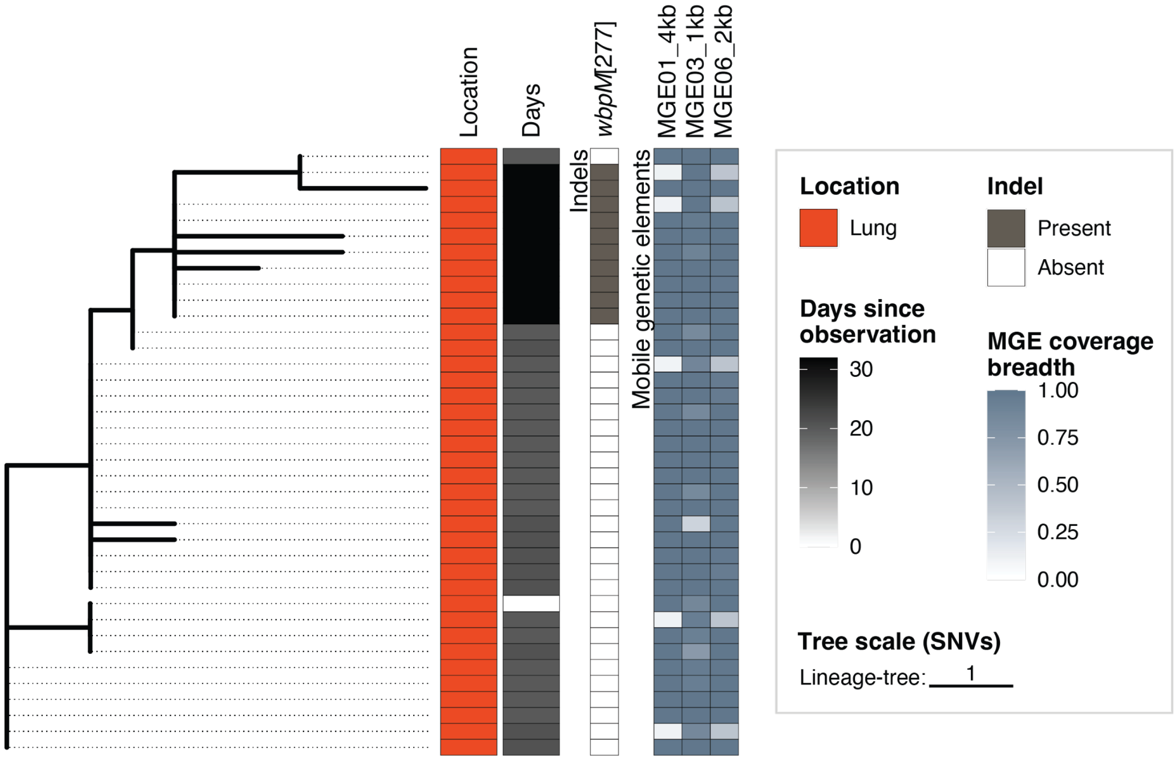


**Figure S10: Population structure of *Pseudomonas aeruginosa* within patient 7.**
Maximum-likelihood phylogeny of the 38 *P. aeruginosa* isolates from patient 7 using the lineage-specific alignment. The tree, based on 13 SNVs, is rooted on GCF_001045685.1 and the tree scale bar represents one SNV. All isolates have been retained from the site of infection (red) and no available sequence type was identified via SRST2. Timepoints since first isolation (in days) are shown in grey scale. Non-singleton indels are shown in grey within a separate heatmap aligned to the tips of the tree with the gene and position of the mutated amino acid in squared brackets. Mobile genetic elements are shown by coverage breadth within a separate heatmap aligned to the tips of the tree if it is supported by at least two isolates (≥ 95% coverage breadth) and lacking in two other isolates (≤ 75% coverage breadth) to reduce complexity. The MGEs are numbered consecutively and the length of the MGE is stated. All identified mutations of *P. aeruginosa* from patient 7 can be found in **Table S7**.

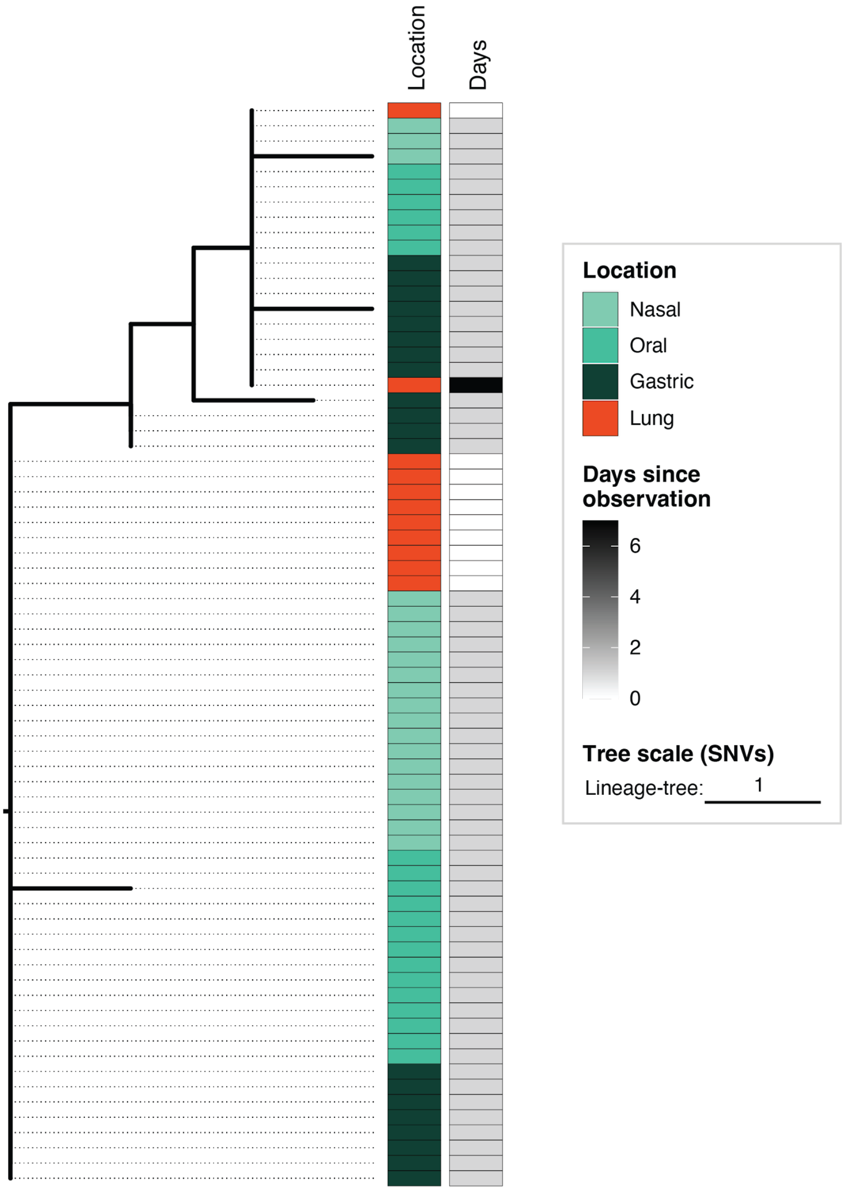


**Figure S11: Population structure of *Klebsiella michiganensis* within patient 10.** Maximum-likelihood phylogeny of the 71 *K. michiganensis* isolates from patient 10 using the lineage-specific alignment. The tree, based on six SNVs, is rooted on GCF_013636415.1 and the tree scale bar represents one SNV. The microbiome isolation source (incl. gastric juice) is shown in shades of green or red for isolates collected from the site of infection. Timepoints since first isolation (in days) are shown in grey scale. No non-singleton indels were identified nor mobile genetic elements supported by at least two isolates (≥ 95% coverage breadth) and lacking in two other isolates (≤ 75% coverage breadth), done to reduce complexity, have been identified. All identified mutations of *K. michiganensis* can be found in **Table S7**.

19

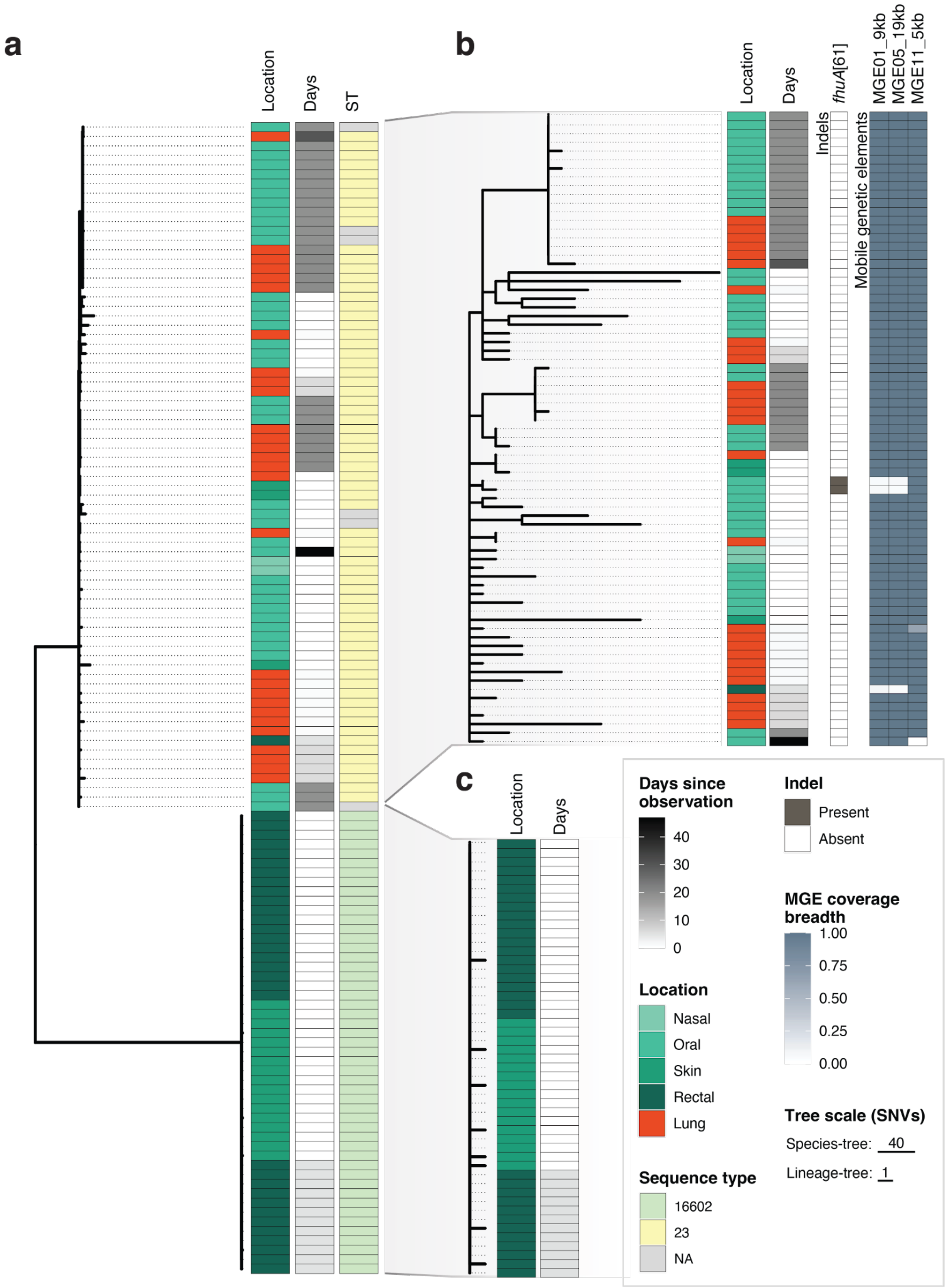


**Figure S12: Population structure of *Escherichia coli* within patient 21.**
**a.** Maximum-likelihood phylogeny of the 122 *E. coli* isolates from patient 21 using the core-genome alignment. The tree is rooted on GCF_000782595.1 and the respective species-specific tree scale bar represents 20 SNV. The microbiome isolation source is shown in shades of green or red for isolates

collected from the site of infection. Timepoints since first isolation (in days) are shown in grey scale. The sequence types (ST) per isolates are aligned to the tips of the tree.

**b-c.** Maximum-likelihood phylogeny per lineage using the lineage-specific alignment. The trees are based on 192 (**b**) and eight (**c**) SNVs, respectively, rooted on GCF_900042785.1 (**b**) and GCF_003856695.1 (**c**), and scaled to the lineage-specific scale bar representing one SNV. Isolation timepoints and source use the same color scheme as in (**a**). For sequence type 21 (**b**), isolates carrying non-singleton indels are shown in grey within a separate heatmap aligned to the tips of the tree with the amino acid location within the gene stated in squared brackets. Mobile genetic elements are shown by coverage breadth within a separate heatmap aligned to the tips of the tree if it is supported by at least two isolates (≥ 95% coverage breadth) and lacking in two other isolates (≤ 75% coverage breadth) to reduce complexity. The MGEs are numbered consecutively and the length of the MGE is stated. No mobile genetic element (≥ 2 isolates ≥ 95% coverage breadth; ≥ 2 isolates ≤ 75% coverage breadth) nor non-singleton indels have been observed for sequence type 16602. All identified mutations of both lineages can be found in **Table S7**.

21

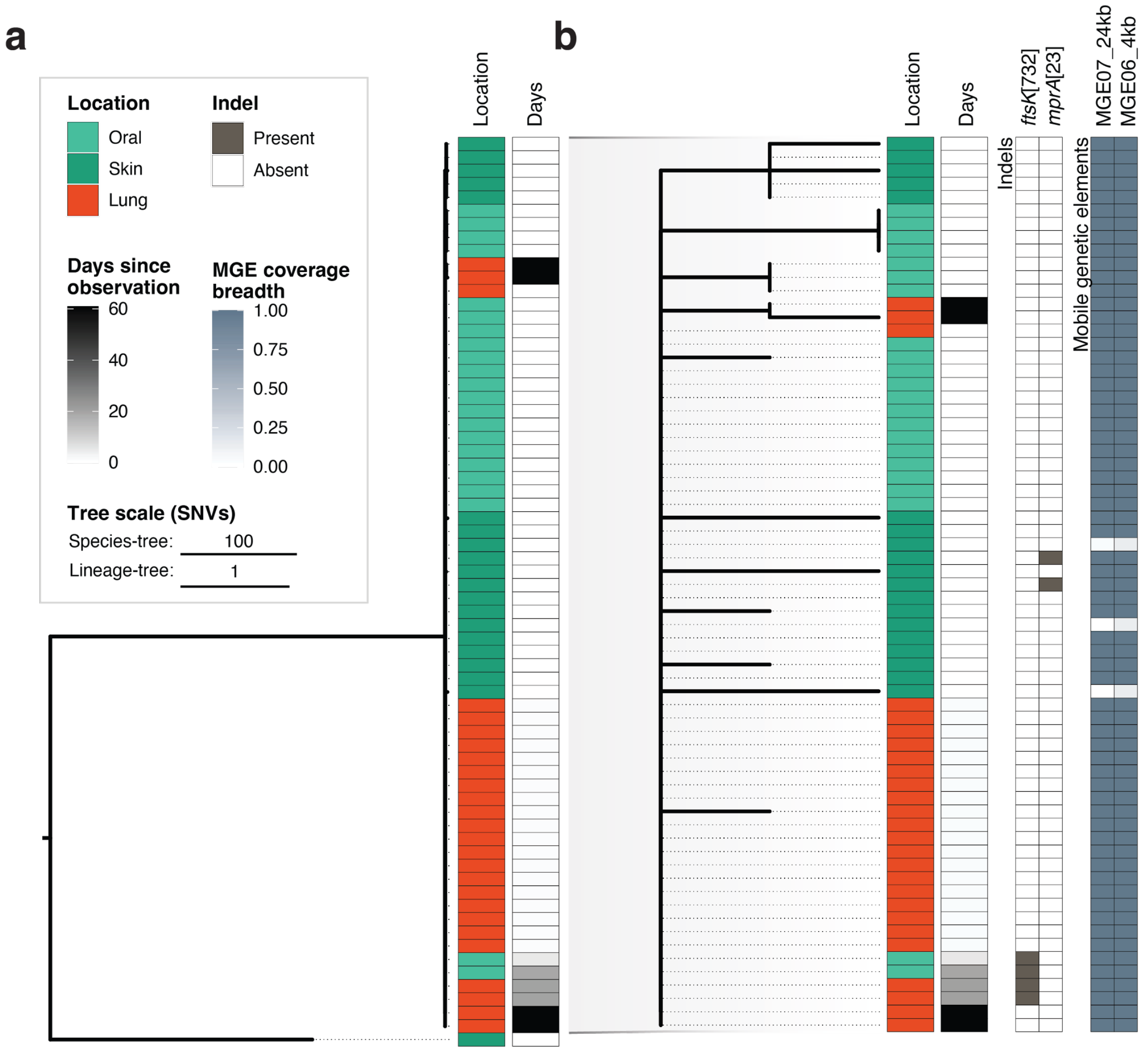


**Figure S13: Population structure of *Proteus mirabilis* within patient 21.**
**a.** Maximum-likelihood phylogeny of the 68 *P. mirabilis* isolates from patient 21 using the core-genome alignment. The tree is rooted on GCF_000069965.1 and the respective species-specific tree scale bar represents 100 SNV. The microbiome isolation source is shown in shades of green or red for isolates collected from the site of infection. Timepoints since first isolation (in days) are shown in grey scale.
**b.** Maximum-likelihood phylogeny of the lineage containing more than one isolate, defined as the major lineage, using the lineage-specific alignment. The tree, based on 18 SNVs, is rooted on GCF_000783875.2 and scaled with the respective lineage-specific scale bar representing one SNV. Isolation timepoints and source use the same color scheme as in (**a**). Non-singleton indels are shown in grey within a separate heatmap aligned to the tips of the tree with the gene and position of the mutated amino acid in squared brackets. Mobile genetic elements are shown by coverage breadth within a separate heatmap aligned to the tips of the tree if it is supported by at least two isolates (≥ 95% coverage breadth) and lacking in two other isolates (≤ 75% coverage breadth). The MGEs are numbered consecutively and the length of the MGE is stated. All identified mutations of *P. mirabilis* can be found in **Table S7**.

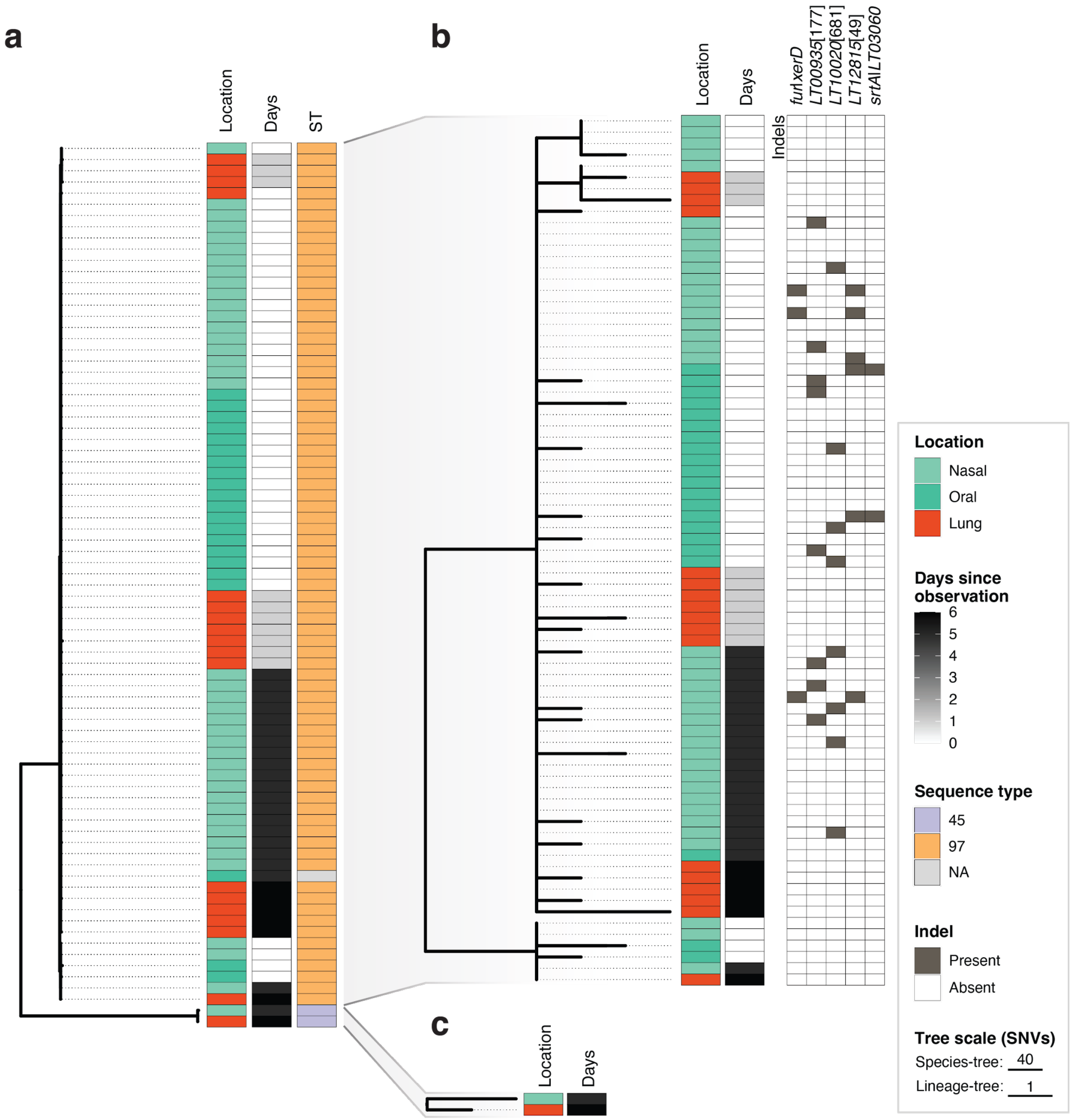


**Figure S14: Population structure of *Staphylococcus aureus* within patient 21.**
**a.** Maximum-likelihood phylogeny of the 79 *S. aureus* isolates from patient 21 using the core-genome alignment. The tree is rooted on GCF_900635305.1 and the respective species-specific tree scale bar represents 40 SNV. The microbiome isolation source is shown in shades of green or red for isolates collected from the site of infection. Timepoints since first isolation (in days) are shown in grey scale. The sequence types (ST) per isolates are aligned to the tips of the tree.
**b-c.** Maximum-likelihood phylogeny per lineage using the lineage-specific alignment. The trees are based on 37 (**b**) and three (**c**) SNVs, respectively, rooted on GCF_900635305.1 (**b**) and GCF_003288395.1 (**c**), and scaled to the lineage-specific scale bar representing one SNV. Isolation timepoints and source use the same color scheme as in (**a**). For sequence type 97 (**b**), non-singleton indels are shown in grey within a separate heatmap aligned to the tips of the tree with the gene and position of the mutated amino acid in squared brackets, while for intergenic indels, adjacent genes are stated, separated by "|". To reduce complexity, mobile genetic elements are filtered to be supported by at least two isolates (≥ 95% coverage breadth) and lacking in two other isolates (≤ 75% coverage breadth) of which none was identified for ST97 (**b**) nor ST45 (**c**). No non-singleton indels were identified for ST45. All identified mutations of both lineages can be found in **Table S7**.

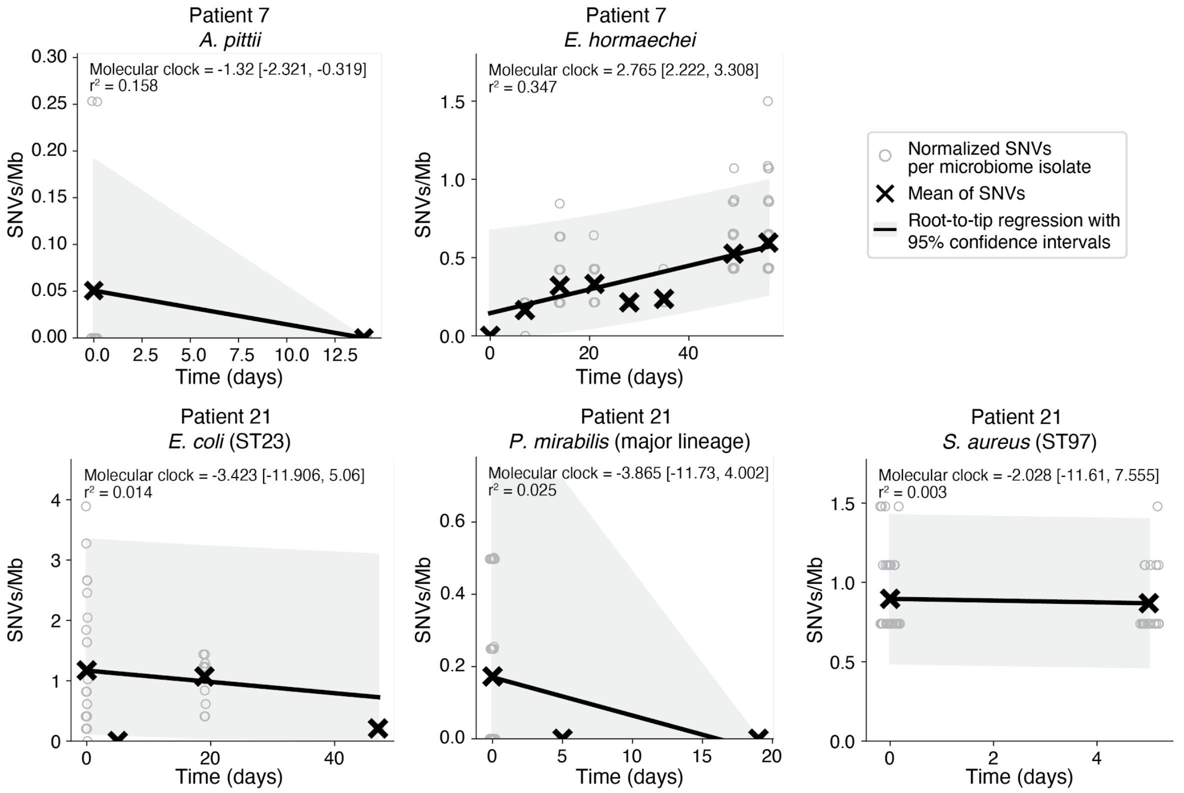


**Figure S15: Inferred molecular clocks from pathogenic lineages.**
Lineage-specific molecular clocks were obtained for all lineages with at least two microbiome-positive samples by fitting a linear regression of the mean normalized SNV counts per megabase (Mb) of all microbiome isolates over time. Individual coverage-breadth normalized SNV counts are shown by circular hollow markers. Regression of mean values (black cross) is depicted by a black line with the 95% confidence intervals in grey. The molecular clock estimates, their 95% confidence intervals, and $r^2$ statistic are shown in the top left of each plot.

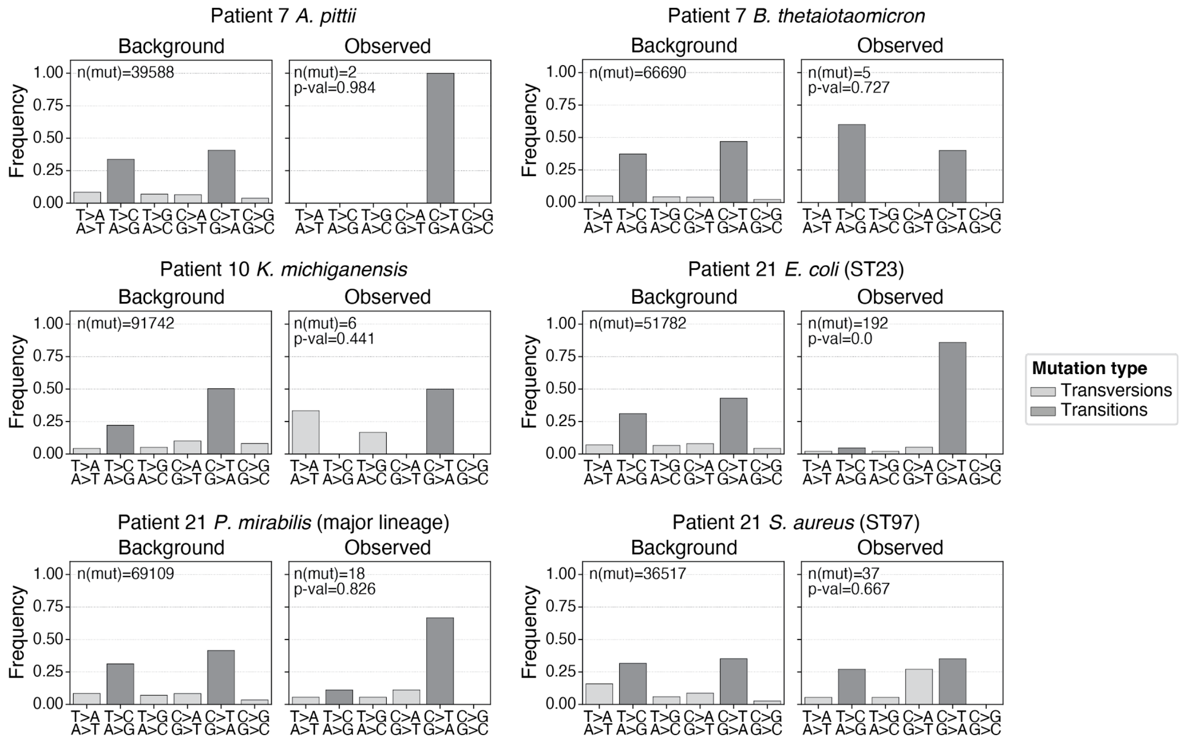


**Figure S16: Mutational spectrum of pathogenic lineages lacking a molecular clock signal.** Observed mutational spectra are shown for all pathogenic lineages detected in the microbiome before or within 6 h of symptom onset with more than eight isolates (prerequisite for TMRCA inference) which lack a molecular clock signal (**Figure S15**). Background frequencies identified across publicly available genomes (**Table S4**) are shown on the left and the observed mutational spectrum on the right per pathogenic lineage. The mutations are categorized within six mutational signatures and color-coded by transitions (light grey) and transversions (dark grey). The number of mutations observed are shown on the top left. Statistical differences ($\chi^2$ test) between background and observed mutational spectra were assessed using transitions and transversions to mitigate the effects of low mutational counts and p-values are shown on the top left in the right panels per lineage.

28

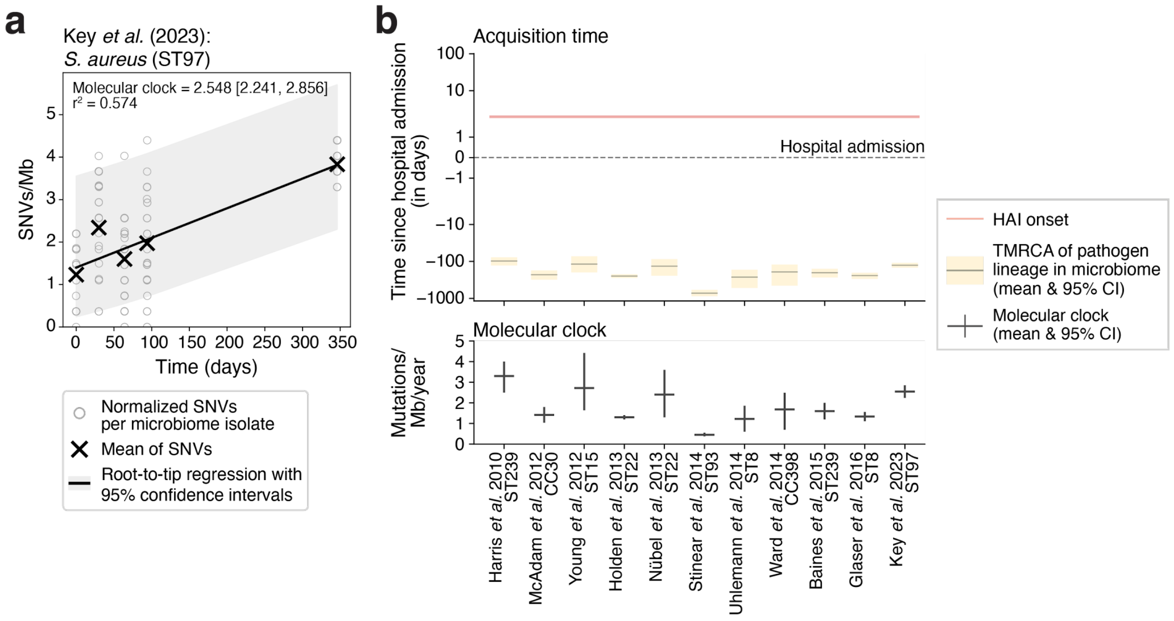


**Figure S17: Molecular clock and TMRCA of alternative published *S. aureus* molecular clocks support inference of *S. aureus* ST97 acquisition before hospitalization.**
**a.** Root-to-tip regression of the reanalysed data from patient 16 of Key *et al.* (2023) [3] utilized for **Figure 2**. Each hollow dot represents the coverage-breadth normalized SNVs counts per megabase (Mb). Linear regression of mean values per timepoint (black cross) is depicted by a black line with the 95% confidence interval shown in grey. The molecular clock estimate is shown on the top left, together with the 95% confidence intervals (in square brackets) and the $r^2$ statistic underneath.
**b.** Inferred time to the most recent common ancestor (upper panel) of *S. aureus* (ST97) in respect to the hospital admission (dashed line) and the HAI onset (red horizontal line) using eleven different published molecular clocks (including Key *et al.* (2023) shown in **Figure 2**) [3–13]. The mean estimate is shown per clock in grey with the 95% confidence intervals shown in yellow. Confidence intervals were calculated using 10,000 bootstraps of the observed genotypes and were jointly combined with random sampling from a gamma distribution fitted to the molecular clocks. The lower panel represents the point estimates of the molecular clocks (horizontal line) and their 95% interval estimates (vertical line) used for the TMRCA inference. The molecular clocks were obtained from eleven published studies with the molecular clock from Key *et al.* (2023) being recalculated following their published thresholds (see **a**).

29

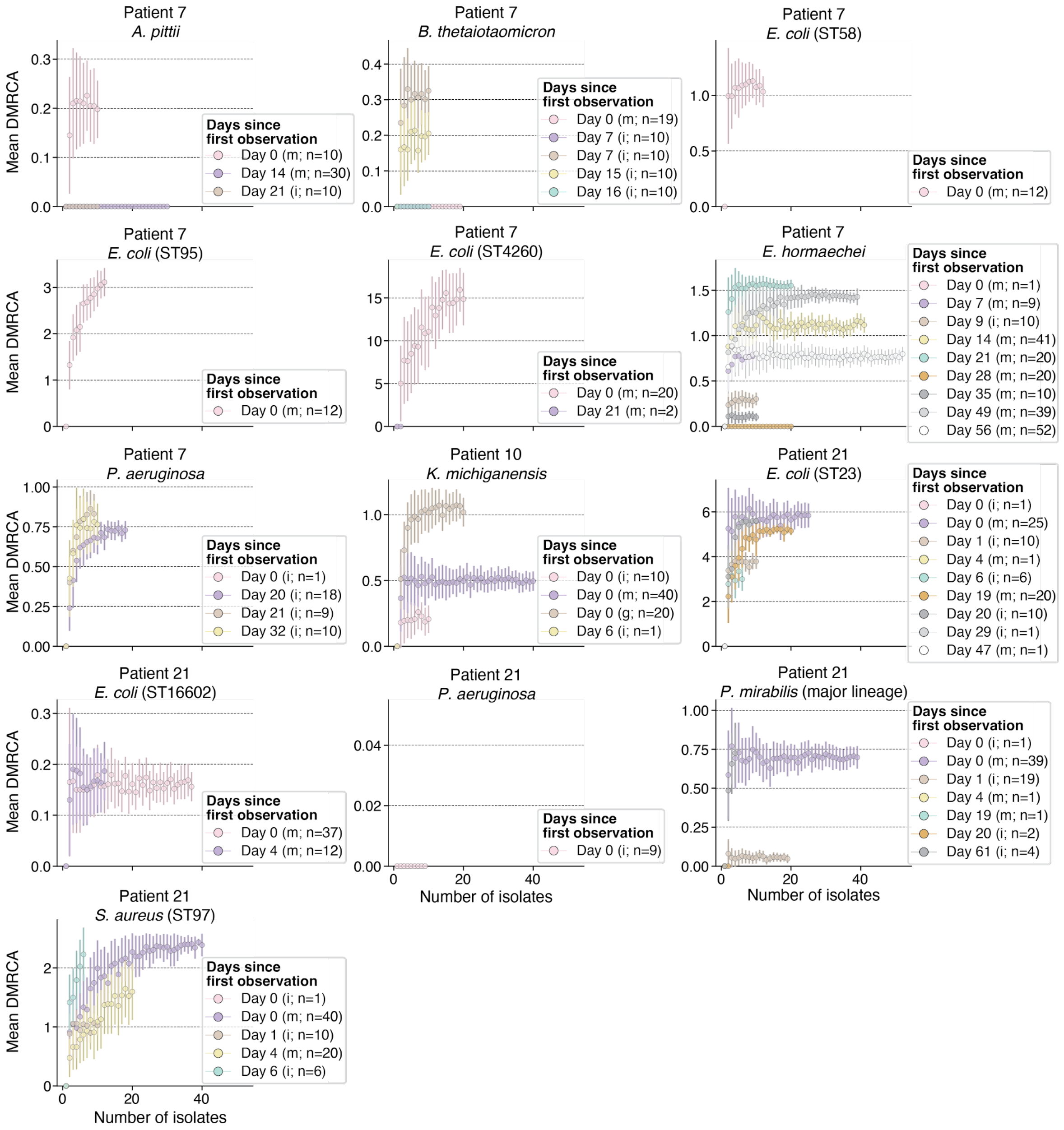


**Figure S18: Body-wide lineage-specific DMRCA collector's curves indicate sufficient sampling for most of the observed lineages per patient and timepoint.**

The DMRCA collector's curves were calculated for all observed lineages with at least five isolates, excluding *E. coli* (ST117) and *E. coli* (ST131) from patient 7, *S. maltophilia* from patient 10, and the minor lineage of *P. mirabilis* and *S. aureus* (ST45) from patient 21. For each distinct timepoint of the retained lineages, 100 random resamples with replacement were generated for every possible subset size $x$ ($1 \leq x \leq n$; $n$ = number of isolates at the respective timepoint). For each resample, the mean DMRCA was calculated. The mean (dot) and the standard error (vertical lines) per subset size $x$ across the 100 resamples is shown. The source of the collected isolates (m = microbiome [collapsed across the four sample sites]; i = site of infection; g = gastric aspirate) is stated in parenthesis in the legend as well as the number of isolates recovered on that timepoint.

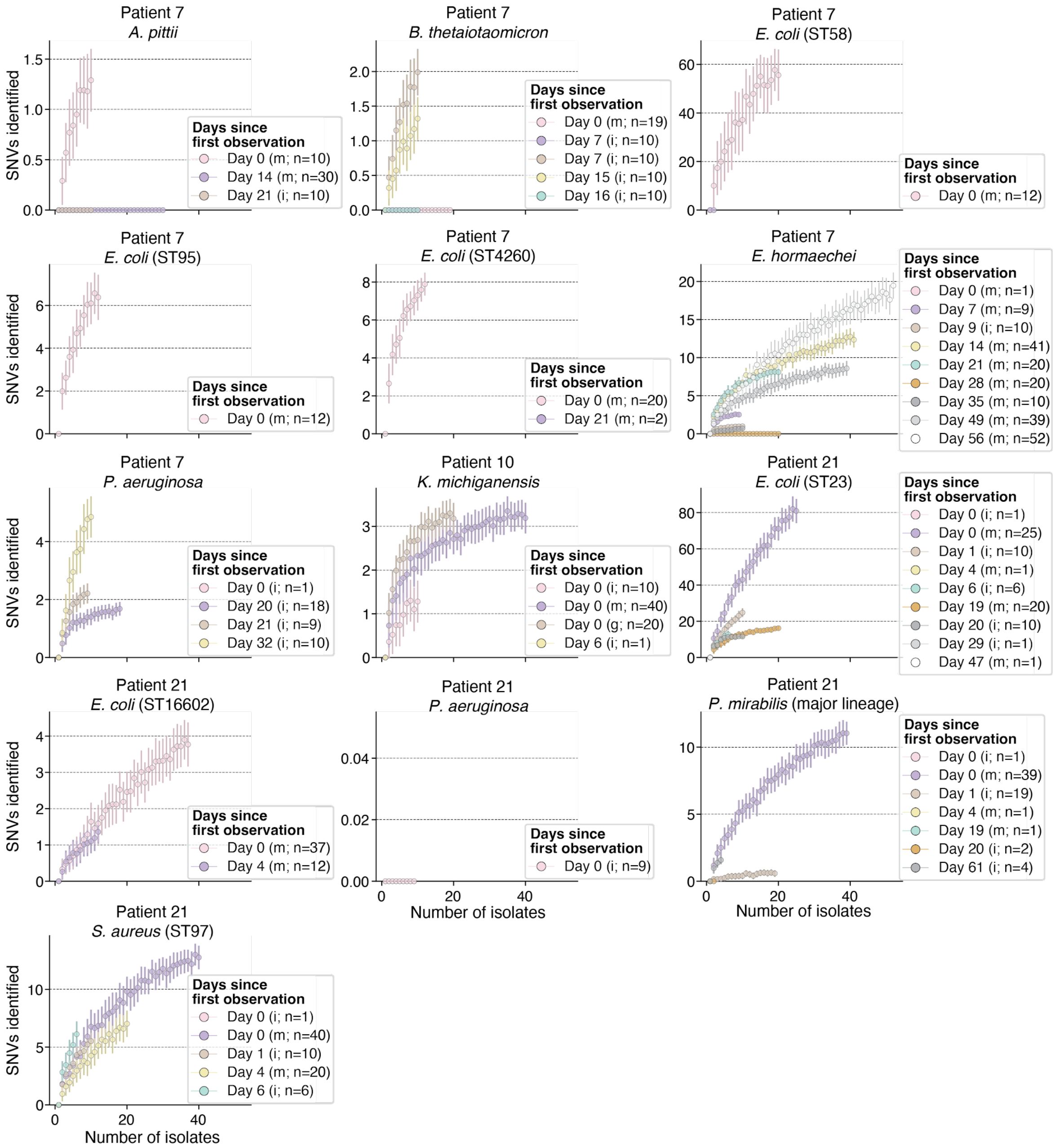


**Figure S19: Body-wide lineage-specific SNV collector's curves suggest unobserved variation within the patient's bodies.**

The SNV collector's curves were calculated for all observed lineages with at least five isolates, excluding *E. coli* (ST117) and *E. coli* (ST131) from patient 7, *S. maltophilia* from patient 10, and the minor lineage of *P. mirabilis* and *S. aureus* (ST45) from patient 21. For each distinct timepoint of the retained lineages, 100 random resamples with replacement were generated for every possible subset size $x$ ($1 \leq x \leq n$; $n$ = number of isolates at the respective timepoint). For each resample, the number of identified SNVs among selected isolates was calculated. The mean (dot) and the standard error (vertical lines) per subset size $x$ across the 100 resamples is shown. The source of the collected isolates (m = microbiome [collapsed across the four samples sites]; i = site of infection; g = gastric aspirate) is stated in parenthesis in the legend as well as the number of isolates recovered on that timepoint.

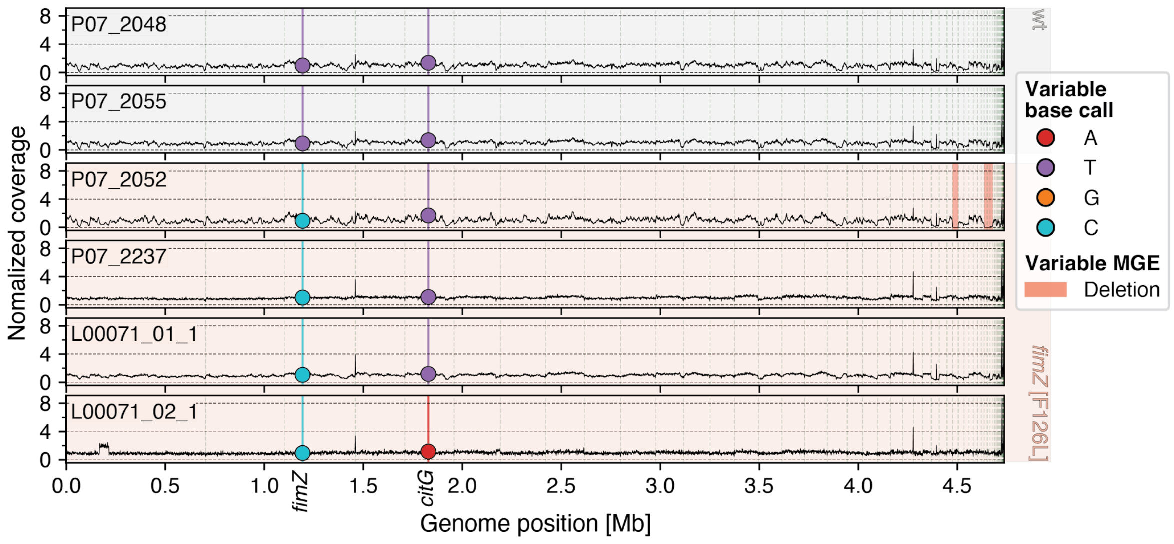


**Figure S20: Genotype comparison of *E. hormaechei* isolates used for *fimZ* [F126L] phenotyping.** Identified genotypes differing among the six isolates used for *fimZ* [F126L] phenotyping are shown with one isolate per panel. The isolate IDs are shown on the top left and the isolates are grouped by genotype with isolates carrying the ancestral, wildtype *fimZ* allele (F126) being on top (P07_2048, P07_2055) shaded in grey, and isolates carrying the derived allele (F126L) are on the bottom (P07_2052, P07_2237 collected from the microbiome; L00071_01_1, L00071_02_1 collected from the site of infection) shaded in salmon. The genome coverage per isolate is shown on the y-axis normalized against the isolate's mean coverage and contig boundaries are indicated by vertical dashed lines. All identified genetic variations (SNV, indel, MGEs) are indicated. For the SNVs, the observed base call is stated at the position. Regions with a deletion are highlighted in red (two deletions in P07_2052) while no indels were identified to deviate among those isolates.

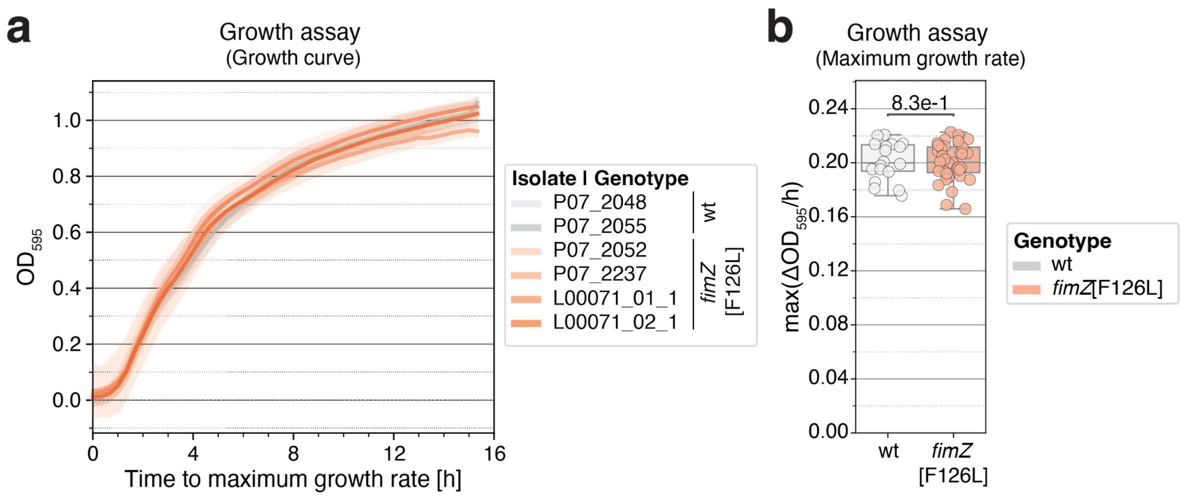


**Figure S21: Growth rate of *E. hormaechei fimZ* [F126L] is similar to the ancestral, wildtype genotype.**

**a.** Growth curves measured by the increase in $OD_{595}$ are shown as the mean (line) and 95% confidence intervals (shadow) per isolate over time. The growth curves are smoothed using a sliding window ± 20 min around each datapoint. Each curve is represented by three independent experimental replicates with three biological replicates per experiment.

**b.** Maximum growth rate identified per genotype across all replicates. Statistical difference was assessed using a Mann-Whitney U test.

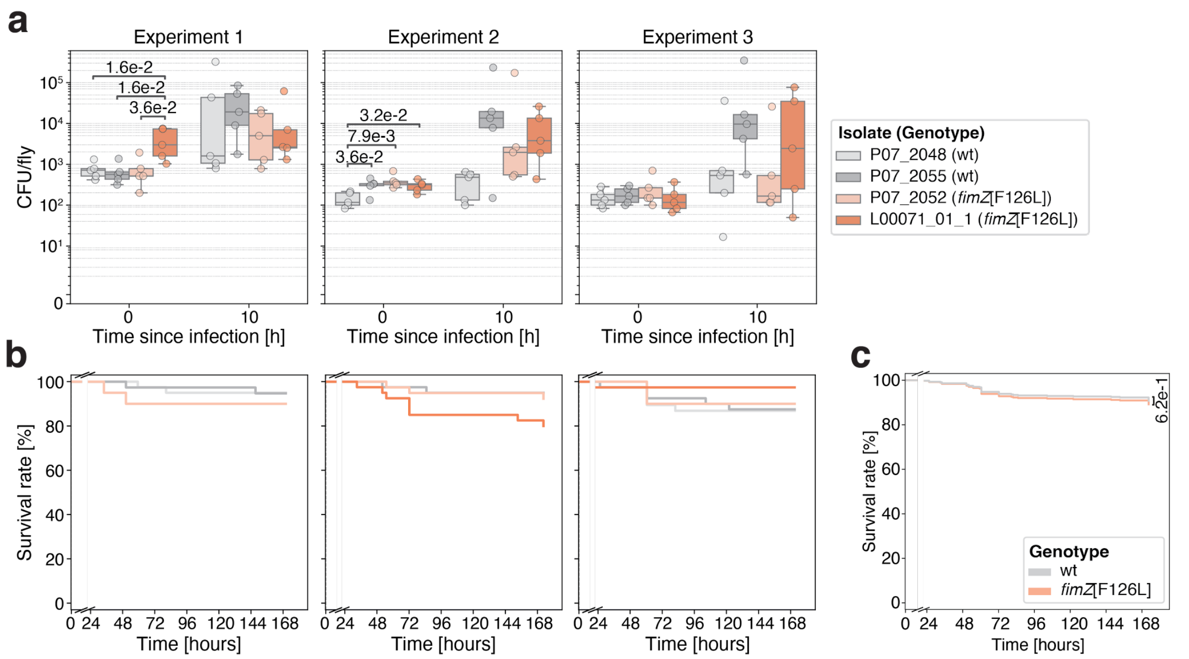


**Figure S22: Systemic infection of *Drosophila melanogaster* DrosDel $w^{1118}$ iso show no difference in virulence of *E. hormaechei fimZ* [F126L] in an immunocompetent host.**

**a.** Colony forming units (CFU) of five single DrosDel $w^{1118}$ iso male flies systemically infected with either the ancestral, wildtype *E. hormaechei* genotype (P07_2048, P07_2055) or the derived *fimZ* [F126L] mutant (P07_2052, L00071_01_1) 0 h and 10 h post infection. Systemic infections have been conducted using a stock solution of $OD_{595}$ 5. Significant differences between isolates per timepoint were assessed using a Mann-Whitney U test.

**b.** Kaplan-Meier curves of 38-40 DrosDel $w^{1118}$ iso male flies per isolate and experiment over one week post infection. Survival curves of isolates with significant differences 0 h post infection (see **a**) have been omitted. The x-axis is broken (8-22 h) to remove the inoculation period.

**c.** Fitted survival curves of DrosDel $w^{1118}$ iso male flies using the Cox proportional hazards model across genotypes (*fimZ* [F126L]: 198 flies; wildtype: 196 flies) using the independent experimental replicates as covariate. The x-axis is broken (8-22 h) to remove the inoculation period. Statistical difference was estimated by fitting a Cox proportional hazards model (p-value = 0.62, Wald test).

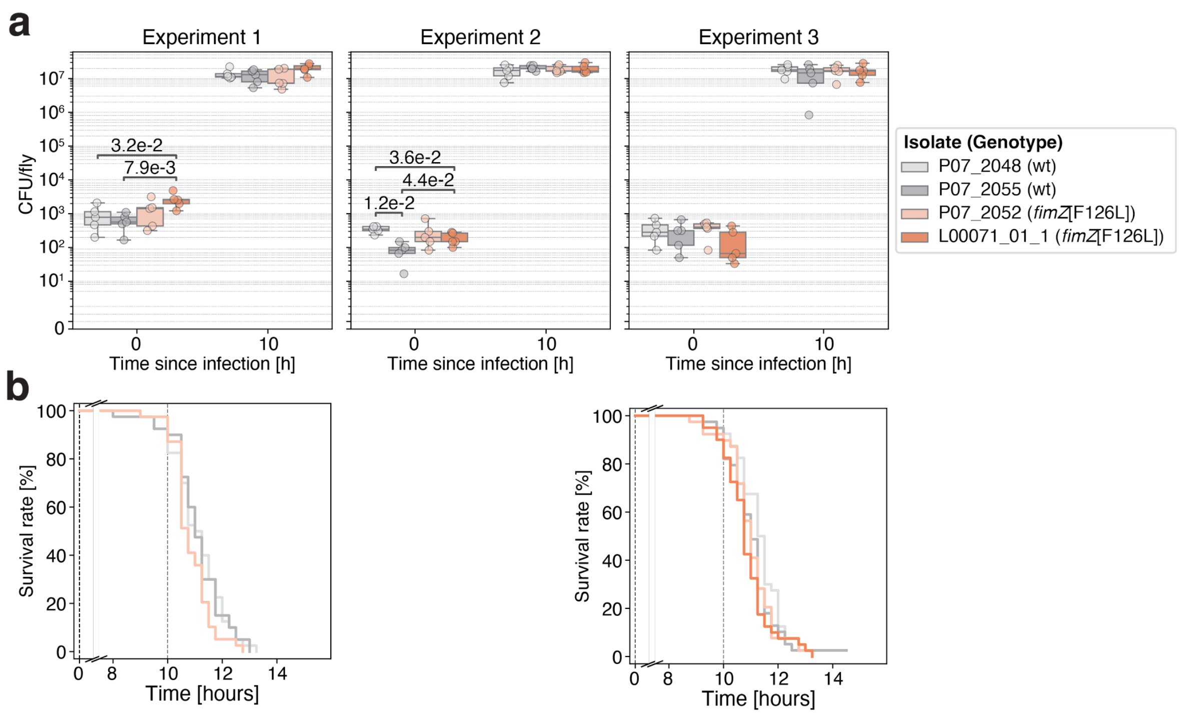


**Figure S23: Systemic infection of *Drosophila melanogaster Relish*$^{E20}$ iso suggests increased virulence of *E. hormaechei fimZ* [F126L] in an immunocompromised host.**

**a.** Colony forming units (CFU) of five single *Relish*$^{E20}$ iso male flies systemically infected with either the ancestral, wildtype *E. hormaechei* genotype (P07_2048, P07_2055) or the derived *fimZ* [F126L] mutant (P07_2052, L00071_01_1) 0 h and 10 h post infection using a stock solution of $OD_{595}$ 5. Significant differences between isolates per timepoint were assessed using a Mann-Whitney U test.

**b.** Kaplan-Meier curves of individual survival experiments using 39-40 flies. Survival curves of isolates with significant differences 0 h post infection (see **a**) have been omitted, resulting in the removal of experiment 2 and L00071_01_1 from experiment 1. The dashed line represents the timepoints of CFU measurements (see **a**). The x-axis is broken (0.5-7.5 h) to remove the inoculation period.

39

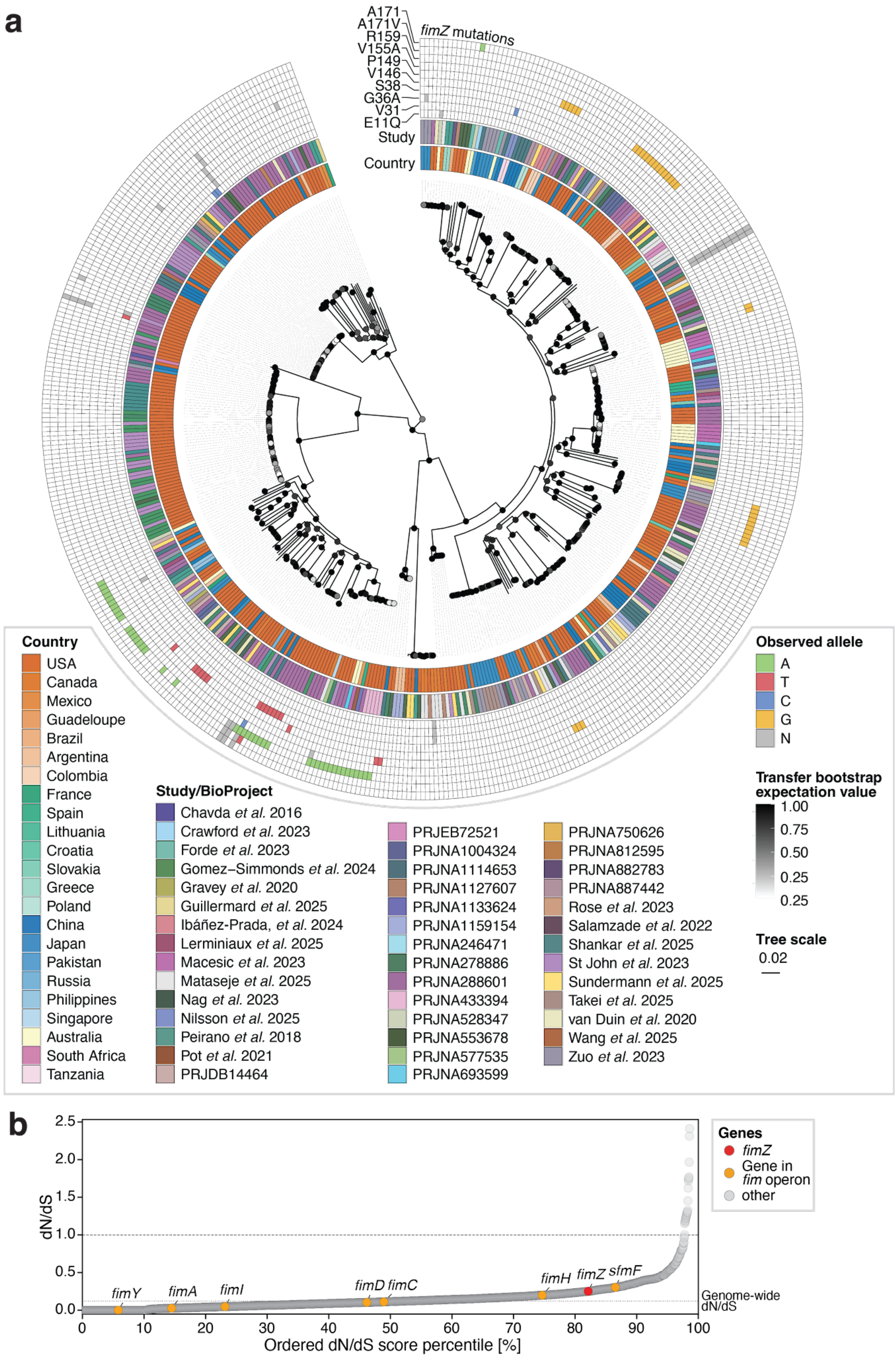

a
A171
A171V
R159
V155A
P149
V146
S38
G36A
V31
E11Q
Study
Country
fimZ mutations
Country
USA
Canada
Mexico
Guadeloupe
Brazil
Argentina
Colombia
France
Spain
Lithuania
Croatia
Slovakia
Greece
Poland
China
Japan
Pakistan
Russia
Philippines
Singapore
Australia
South Africa
Tanzania
Study/BioProject
Chavda et al. 2016
Crawford et al. 2023
Forde et al. 2023
Gomez–Simmonds et al. 2024
Gravey et al. 2020
Guillermard et al. 2025
Ibáñez-Prada, et al. 2024
Lerminiaux et al. 2025
Macesic et al. 2023
Mataseje et al. 2025
Nag et al. 2023
Nilsson et al. 2025
Peirano et al. 2018
Pot et al. 2021
PRJDB14464
PRJEB72521
PRJNA1004324
PRJNA1114653
PRJNA1127607
PRJNA1133624
PRJNA1159154
PRJNA246471
PRJNA278886
PRJNA288601
PRJNA433394
PRJNA528347
PRJNA553678
PRJNA577535
PRJNA693599
PRJNA750626
PRJNA812595
PRJNA882783
PRJNA887442
Rose et al. 2023
Salamzade et al. 2022
Shankar et al. 2025
St John et al. 2023
Sundermann et al. 2025
Takei et al. 2025
van Duin et al. 2020
Wang et al. 2025
Zuo et al. 2023
Observed allele
A
T
C
G
N
Transfer bootstrap expectation value
1.00
0.75
0.50
0.25
0.25
Tree scale
0.02
b
dN/dS
2.5
2.0
1.5
1.0
0.5
0.0
fimY
fimA
fimI
fimD
fimC
fimH
fimZ
sfmF
Genome-wide dN/dS
0
10
20
30
40
50
60
70
80
90
100
Ordered dN/dS score percentile [%]
Genes
fimZ
Gene in fim operon
other

**Figure S24: Analysis of 480 publicly available *E. hormaechei* whole-genome sequences reveals the *fim* operon, including *fimZ*, to evolve under purifying selection in the clinical context.**

**a.** Maximum-likelihood tree of 480 publicly available *E. hormaechei* whole-genome sequences (79,610 SNVs) used to evaluate the frequency of non-synonymous *fimZ* mutations in the respiratory tract. Transfer bootstrap expectation (TBE) values of 1,000 bootstraps are marked on the respective nodes via grey scale. The whole-genome sequences were obtained from the Pathogen Detection database [1] and the CDC HAI-Seq Gram-negative bacteria BioProject [2]. The original studies, if identified, or their BioProject IDs as well as the country of isolation for each genome are shown as heatmaps on the two inner circles. The outer heatmap shows all observed SNVs in *fimZ* annotated by the amino acid change. The heatmap is colored by observed alternative alleles (green: A; red: T; blue: C; yellow: G) with the reference call shown in white and ambiguous calls (N) in grey. Synonymous mutations V146, P149, and R159 show dispersed phylogenetic placement likely reflecting topological uncertainties arising from recombination; however, the overall species' population structure is robustly resolved as shown by the TBE values.

**b.** Genome-wide evaluation of the gene-specific dN/dS scores for all mutated genes (n = 2,909). dN/dS scores were ordered and each gene is represented by a grey circle. Genes with non-synonymous mutations only (n = 41; division by 0 and therefore unable to derive a dN/dS score) are not shown and genes with a dN/dS score of 0 (n = 307) were ordered by gene length. Type I fimbriae genes part of the *fim* operon were highlighted in orange with *fimZ* highlighted in red and their respective gene name stated above. Neutral expectation (dN/dS = 1) is marked by a horizontal dashed line and the genome-wide dN/dS score (0.123) is indicated by a dotted line.

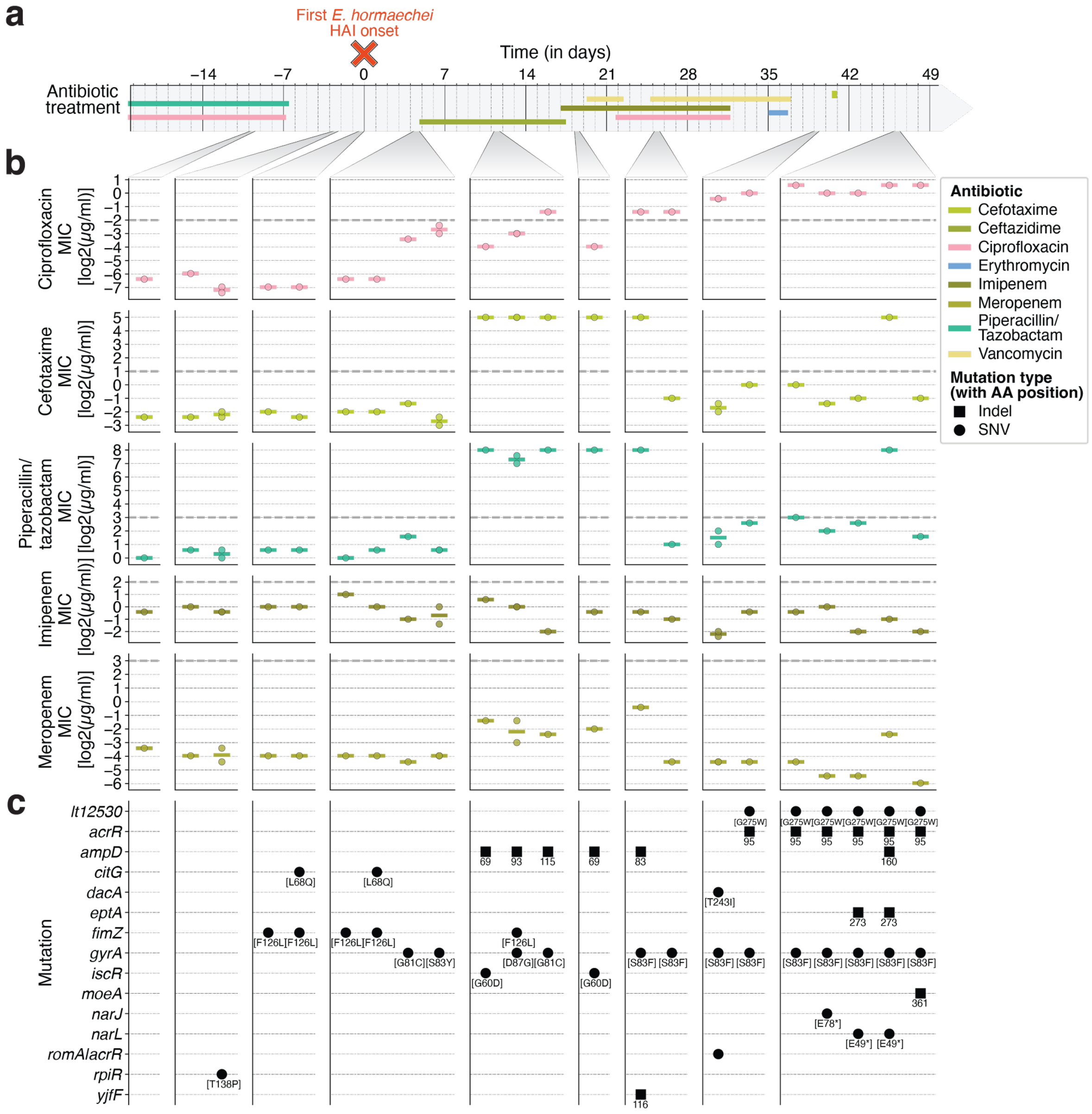


**Figure S25: Minimal inhibitory concentrations of *Enterobacter hormaechei* isolates against antibiotics administered to patient 7 during hospital stay.**

**a.** Antibiotic treatment episodes of patient 7 twenty days prior to the first observation of *E. hormaechei* until three days after the last microbiome sample of patient 7. First HAI onset by *E. hormaechei* within patient 7 marked above with a red cross.

**b.** Minimal inhibitory concentration of selected isolates representing the major phylogenetic sub-clades (**Figure S9**) against all tested antibiotics (1x fluoroquinolone [ciprofloxacin], and 4x beta-lactam [cefotaxime, imipenem, meropenem, piperacillin/tazobactam]). Each column represents a sampled timepoint aligned to (**a**) with the tested genotype. Each circular marker represents one isolate and the line represents the mean of the minimum inhibitory concentration (MIC) per genotype. Clinical breakpoints defined by the EUCAST Clinical Breakpoints Tables (v15.0) are shown by grey dashed lines.

**c.** SNVs (including the *fimZ* [F126L] mutation) and indels for each individual genotype are shown. Below each SNV the position and amino acid change is shown and for indels the amino acid start position is shown. One intergenic mutation, a SNV, located between divergently transcribed genes has been identified (*romA*|*acrR*): 68 nt downstream of *romA* and 124 nt upstream of *acrR*.
